# A predisposing effect of HLA class II genes in celiac disease by skewing the naïve CD4^+^ T-cell receptor repertoire

**DOI:** 10.1101/2025.07.15.663863

**Authors:** Ida Lindeman, Aengus Officer, Shiva Dahal-Koirala, Louise F. Risnes, Rebecka Hjort, Knut E. A. Lundin, Eivind Ness-Jensen, Corey T. Watson, Ludvig M. Sollid

**Affiliations:** Norwegian Coeliac Disease Research Centre, University of Oslo, Oslo, Norway; Department of Immunology, Oslo University Hospital - Rikshospitalet, Oslo, Norway; Institute of Clinical Medicine, University of Oslo, Oslo, Norway; Department of Gastroenterology, Oslo University Hospital - Rikshospitalet, Oslo, Norway; HUNT Research Centre, Department of Public Health and Nursing, NTNU, Norwegian University of Science and Technology, Levanger, Norway; HUNT Center for Molecular and Clinical Epidemiology, Department of Public Health and Nursing, NTNU, Norwegian University of Science and Technology, Trondheim, Norway; Department of Medicine, Levanger Hospital, Nord-Trøndelag Hospital Trust, Levanger, Norway; Upper Gastrointestinal Surgery, Department of Molecular Medicine and Surgery, Karolinska Institutet and Karolinska University Hospital, Stockholm, Sweden; Department of Biochemistry and Molecular Genetics, School of Medicine, University of Louisville, KY, USA

## Abstract

Polymorphisms of human leukocyte antigen (HLA) genes confer risks for many human diseases. For predisposing effects relating to T-cell receptor (TCR) recognition of peptide-HLA, the effect can be both selection of the TCR repertoire and preferential presentation of disease-driving epitopes. In celiac disease (CeD) HLA-DQ2.5 predisposes by presenting deamidated gluten peptides to CD4^+^ T cells that typically employ stereotyped TCRs. Here we analyzed whether genetic variants within the HLA and TR loci shape the naïve TCR repertoire. We sequenced the αβ TCR repertoires of naïve CD4^+^ T cells of 103 CeD subjects and 103 controls and performed gene usage quantitative trait loci analyses. The naïve CD4^+^ TCR repertoire was significantly affected by HLA and TRA and TRB polymorphisms. Individuals carrying the HLA-DQ2.5 allotype exhibited increased frequencies of TCR genes involved in stereotyped recognition of gluten epitopes thus demonstrating a disease-predisposing effect of HLA by selection of a disease-relevant TCR repertoire.

## Introduction

Autoimmune conditions develop as a result of interplay between genes and environmental factors^1^. The gene-enviroment interactions result in tissue damaging immune responses that mainly involve the T cells and B cells of the adaptive immune system. A central component of the adaptive immune system is major histocompatibility complex (MHC) molecules. These molecules, termed human leukocyte antigen (HLA) molecules in man, bind and present antigenic peptide fragments to T cells. HLA class I (i.e. A, B or C) molecules present antigen to CD8^+^ T cells whereas HLA class II (i.e. DR, DQ or DP) molecules present antigen to CD4^+^ T cells. Both CD8^+^ T cells and CD4^+^ T cells use an αβ T-cell receptor (TCR) to recognize the peptide-HLA (pHLA) complex. The αβ TCR is generated through the process of somatic recombination of variable (V), diversity (D) and joining (J) gene segments during T-cell development in the thymus. In the thymus the T cells also undergo positive and negative selection leading to formation naïve cells that can encounter antigen in the periphery. The TCR repertoire of naïve T cells is vast allowing recognition of many antigens. Both the HLA and TCR genes display high degrees of genetic variation at the population level which could confer influence on the development of disease.

The genetic influence of HLA for autoimmune diseases has been known for six decades^2^, and more recent genome wide association studies (GWAS) have confirmed the principal role of HLA genes in the etiology of these diseases^3^. Conceivably, predisposing effects of HLA that relate to TCR recognition can act by preferential presentation of peptide antigens to pathogenic T cells in the periphery (peripheral hypothesis), or they can work by affecting selection of T cells during their maturation in thymus resulting in a biased representation of potentially pathogenic T cells (central hypothesis)^4,5^. The central and peripheral hypotheses are not mutually exclusive. For many diseases the insight into which of these mechanisms are operating is limited due to the lack of knowledge about the antigens that drive immune reactions causing the disease. In this regard CeD is an exception as in this disease the target antigens (gluten peptides and transglutaminase 2) are known and the HLA association is well-described. Virtually all CeD individuals express HLA-DQ2.5, HLA-DQ2.2 or HLA-DQ8; allotypes that uniquely can bind gluten peptides that have been post-translationally deamidated by transglutaminase 2, thereby allowing efficient presentation of gluten antigen to CD4^+^ T cells^6–8^. Interestingly, T cells isolated from different CeD patients that recognize the same epitope in the context of a given HLA-DQ molecule often use similar TCRs encoded by a restricted set of α and β V genes (TRAV and TRBV), indicating that the T-cell response to gluten in CeD is stereotypical^9^.

HLA variants could influence the composition of the TCR repertoire at different stages of the lifespan of a T cell, as both thymic selection and clonal expansion of antigen-experienced T cells require interactions between the TCR and pHLA complexes. Previous studies have shown that HLA influences the expressed TCR repertoire^10,11^, but were unable to conclusively establish a role during thymic selection. These studies were based on bulk sequencing of a mix of antigen-experienced and naïve T cells^10,11^, making it hard to establish how much of the HLA-effect is due to clonal expansion versus thymic selection. Another study focused on umbilical cord blood, which consists mainly of naïve T cells, and demonstrated the presence of contacts between TCRβ-chain and HLA residues leading to biased TRBV gene usage^12^. While highly valuable, the abovementioned studies did not differentiate between CD4^+^ and CD8^+^ T cells, potentially confounding the analyses as the two populations exhibit distinct TCR biases, and may play differential roles in disease pathogenesis^13^. Some additional evidence in favor of the central hypothesis comes from the finding that HLA risk alleles restrict the hypervariable complementarity determining region 3 (CDR3) regions in naïve TCR repertoires in the direction of disease-relevant CDR3 sequences^14^, and that the HLA alleles of an individual can be inferred from the antigen-experienced TCR repertoire^15^.

Genetic variation within the TR loci has also been reported to influence the composition of the TCR repertoire^10^. The observed biases in TCR repertoires associated with various autoimmune diseases^16^ and the critical functional links between the HLA and TCR make TR loci plausible disease susceptibility candidates, particularly in diseases for which HLA risk alleles play a prominent role. However, with the exception of TCR genes predisposing to narcolepsy^17,18^, the TR loci have been largely overlooked by GWAS due to a poor SNP coverage in the genotyping array most commonly used for immune-mediated diseases^19^.

In the present study we have assessed the effect of HLA allotypes, as well as genetic variants within the TR loci (i.e. TRA and TRB) themselves, on the peripheral blood naïve CD4^+^ TCR repertoire of 206 subjects, of which half had been diagnosed with CeD. By eliminating the confounding presence of antigen-experienced or CD8^+^ T cells, we demonstrate strong HLA-dependent effects on the naïve CD4^+^ TCR repertoire, especially for TRA, indicating that HLA-DQ allotypes predispose to CeD by central effects influencing the selection of naïve T cells in the thymus along with the previously known peripheral effects affecting presentation of gluten peptides to T cells in the gut. Additionally, we identify *cis*-effects for several TRA and in particular TRB genes. Thus, both variants of HLA and TCR genes influence the composition of the naïve CD4^+^ TCR repertoires and may in this manner predispose to immune conditions involving CD4^+^ T cells.

## Results

### Generation of naïve CD4^+^ T cell AIRR-seq libraries, SNP genotyping and HLA typing

We sampled and cryopreserved peripheral blood mononuclear cells (PBMCs) from 103 CeD cases and 103 controls from the fourth population-based Trøndelag Health Study (HUNT4) in Norway^20–22^ (Fig. 1a). Demographic details of the study subjects are shown in Fig. 1b,c. From the PBMCs we isolated RNA from FACS-sorted naïve CD4^+^ T cells (Fig. 1a and Supplementary Fig. 1a) and generated adaptive immune receptor repertoire sequencing (AIRR-seq) libraries for all 206 individuals. To limit technical biases, we aimed for a balanced concentration of RNA template and sequencing reads for controls and CeD subjects, resulting in similar numbers of sequencing reads, unique molecular identifiers (UMIs) and unique TRA and TRB sequences detected for the two subject groups (Fig. 1d,e and Supplementary Fig. 1b,c).

**Figure 1.**
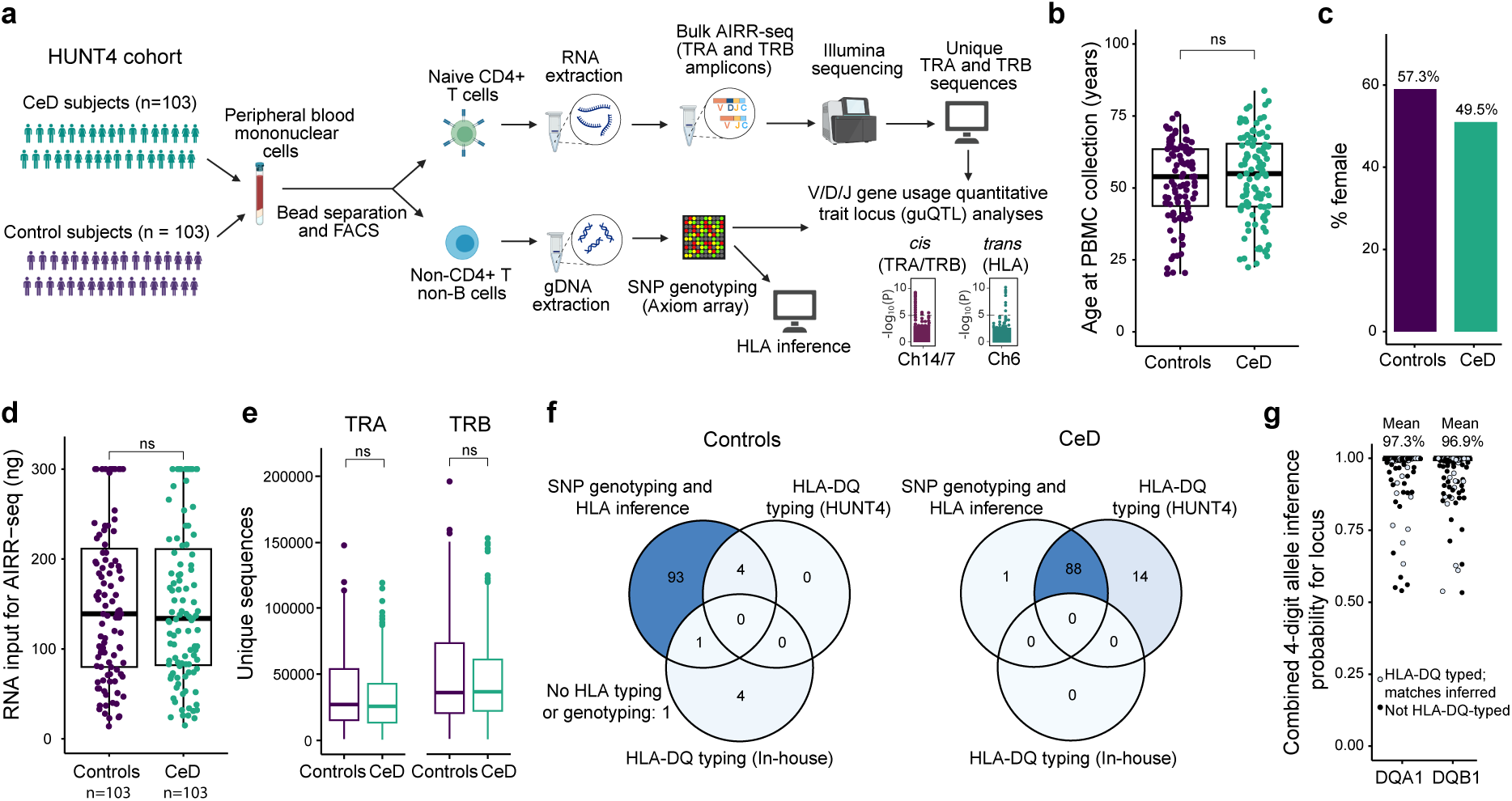
Approach, sample cohort characteristics and sequencing statistics. (**a**) An illustration of our overall approach. (**b,c**) Cohort characteristics of control and CeD subjects, showing age at time of sample collection (**b**) and biological sex distribution (**c**). (**d,e**) AIRR-seq metrics for naïve CD4^+^ T cells isolated from peripheral blood of control and CeD subjects, showing RNA concentration used for preparation of sequencing libraries (**d**) and number of unique TRA and TRB sequences after processing (**e**). (**f**) Genotyping and HLA typing approach for control and CeD subjects. (**g**) HLA allele inference probability reported by the Axiom HLA Analysis tool. Statistical significance was calculated with a Wilcoxon rank sum test (**b**,**d,e**).

We also isolated genomic DNA from most of the individuals, genotyped single nucleotide polymorphisms (SNPs) using an Axiom Human Genotyping SARS-CoV-2 Research Array, and performed quality control and manual inspection of potential contamination (Supplementary Fig. 2). Genotyped SNPs were subsequently used for HLA inference (Fig. 1f,g). We additionally obtained sequencing-based HLA-DQ typing data for four of the controls and all but one of the CeD subjects (results reported elsewhere in Hjort *et. al*, submitted), and determined the HLA-DQ type in-house by qPCR for five of the controls for which no SNP genotyping data was available or the HLA inference failed (Fig. 1f). The HLA inference results matched the HLA-DQ allotype obtained by sequencing or qPCR in all cases where multiple methods had been applied for the same subjects (Fig. 1g). From the SNPs passing quality control and filtering steps, we extracted SNPs on chromosome 6 as well as SNPs within 1 million bases (1Mb) of the start and end positions of the TRA locus on chromosome 14 and TRB locus on chromosome 7 for gene usage quantitative trait locus (guQTL) analyses.

### Genetic variants in the TR loci affect gene usage frequencies in the naïve TCR repertoire

In order to determine whether variation in the naïve CD4^+^ TCR repertoire is influenced by genetic polymorphisms within the TR loci, we focused our analysis on control subjects to rule out disease-specific effects. We first assessed if genetic variants within the TRA and TRB loci influence the expression of individual V, D or J genes by a *cis*-guQTL approach. Using a linear regression framework, we identified a total of 39/685 and 70/262 tested SNPs that were associated with at least one TRA or TRB gene, respectively, after Bonferroni correction (*P* = 7.3 × 10^-5^ (TRA), *P* = 1.9 × 10^-4^ (TRB); Fig. 2a); we refer to these SNPs as guQTLs. The vast majority of the guQTLs were located within the TRA or TRB loci, rather than within the 1 Mb included at each end of the two loci.

**Figure 2.**
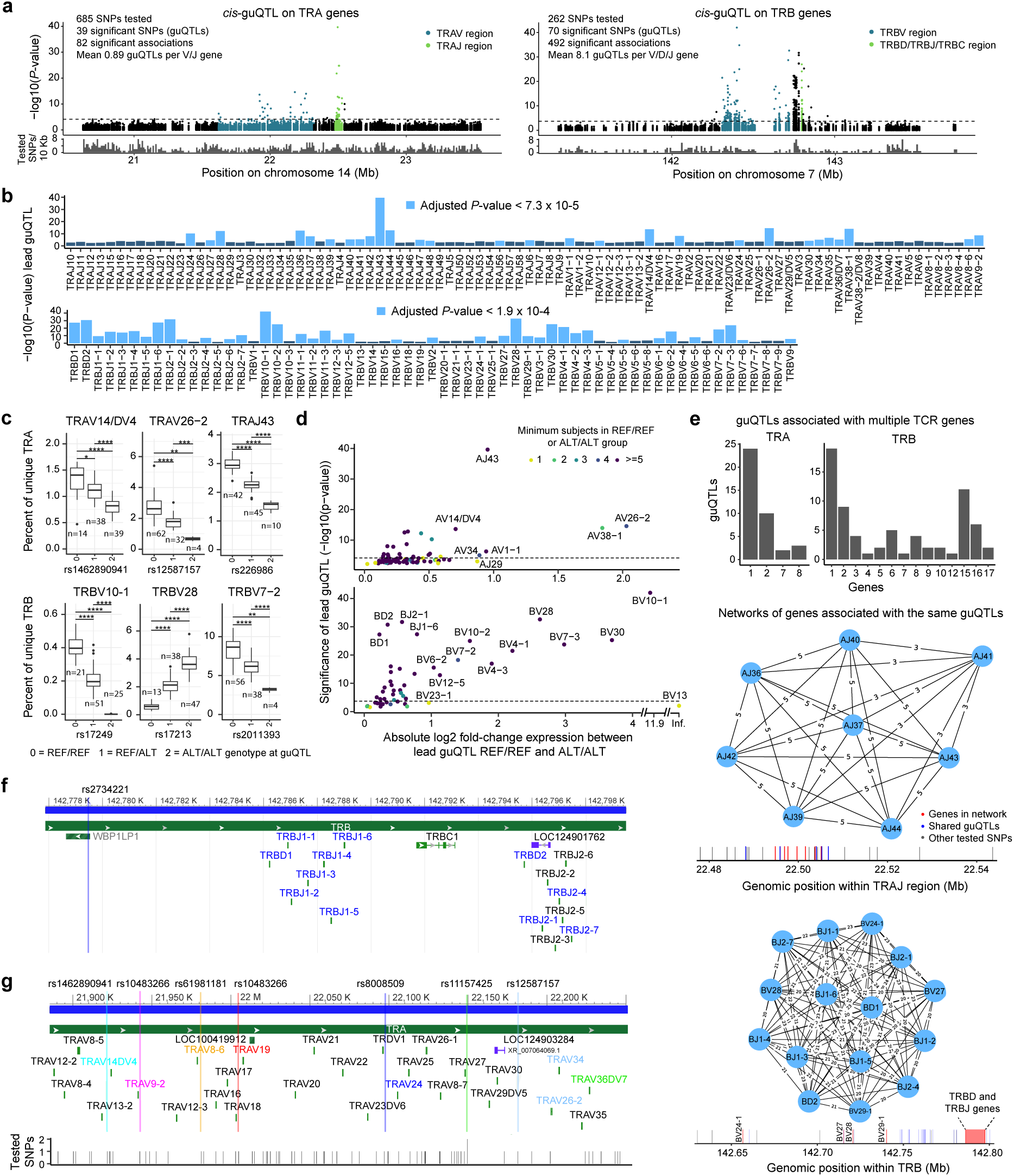
*Cis*-guQTL analysis of TRA and TRB gene usage in naïve CD4^+^ T cells of control subjects. (**a**) Manhattan plots showing SNP associations across V, D and J genes within 1 Mb of the TRA (left) and TRB (right) loci. Dashed lines indicate Bonferroni-corrected significance thresholds. Lower tracks show tested SNPs/10 Kb. (**b**) Significance of lead *cis*-guQTLs for each gene. (**c**) Examples of *cis*-mediated effects of lead guQTL genotype on gene expression. (**d**) Fold-change in expression between subjects homozygous for the REF or ALT lead guQTL variant for each gene. (**e**) guQTLs associated with one or more TRA or TRB gene (upper) and biggest networks of TRA (middle) and TRB (lower) genes associated with the same guQTLs. Gene tracks indicate positions of guQTLs, affected genes and tested SNPs for each network. (**f,g**) Examples of location of lead guQTLs with respect to their affected TRB (**f**) or TRA (**g**) genes.

All gene types (V, D and J) were among those associated guQTLs, with 13/42 TRAV, 15/50 TRAJ, 26/46 TRBV, 2/2 TRBD and 11/13 TRBJ genes showing significant associations with at least one guQTL (Fig. 2b and Supplementary Fig. 3). For some of these genes, the gene usage frequency differed by >8-fold between genotypes (Fig. 2c,d). For example, individuals homozygous for the alternative variant (ALT/ALT) of rs17249 showed virtually no expression of TRBV10-1, while subjects heterozygous (REF/ALT) or homozygous for the reference variant (REF/REF) had significantly higher usage frequency (Fig. 2c). rs17249 is located within the exon sequence of TRBV10-1 and has previously been shown to lead to loss of function due to the introduction of a stop codon^23,24^. Another gene, TRBV28, which was associated with rs17213, showed marked differential usage between genotypes but was observed at >0% frequency in all individuals. Notably, rs17213 is found within the spacer region of the recombination signal sequence (RSS) of TRBV28, and has previously been shown to affect the level of TRBV28 expression in total PBMCs^11^. This result confirms that germline variation can impact usage frequencies of TRBV28 alleles that only differ in their RSS sequence^25^.

Many of the guQTLs were associated with more than one TRA or TRB gene, and we identified networks of genes being affected by the same set of guQTLs (Fig. 2e). These genes were located in close genomic proximity (Fig. 2f and Supplementary Fig. 3c), suggesting that they may be regulated by the same molecular factor(s) or that the guQTLs form linkage disequilibrium (LD) blocks. However, it is possible that incomplete SNP coverage within the TRA and TRB loci could limit our ability to identify guQTLs and affected genes; for instance, only one SNP was present near the TRBD/J region (Fig. 2f), and there are potentially varying degrees of LD within the TRA and TRB loci^26–28^. Finally, many of the V genes had their lead guQTL located within close proximity to the gene segment (Fig. 2g), indicating that these genes may be regulated by variants in or near the genes themselves, for instance in their RSSs or leader sequences.

### The HLA class II region exerts control over naïve TCR repertoires by affecting the frequency of in particular TRA genes

We next performed a *trans*-guQTL analysis aimed at HLA, first analyzing 24690 SNPs located across the entire chromosome 6. Almost all significant associations to the individual TRA or TRB genes could be mapped to the HLA region (Fig. 3a, left panel). Notably, the association was overall stronger for TRA genes compared to TRB genes, with respect to total guQTLs, mean guQTLs per gene and mean *P* value of lead guQTLs. We then limited our analysis to the 6791 SNPs within 1 Mb of the beginning and end of the HLA region, and found that the most significant guQTLs were located in the HLA class II region (Fig. 3a, right panel). Our *trans*-guQTL analysis found 25/92 (27%) TRA genes (17 V genes and 8 J genes) and 11/61 (18%) TRB genes (11 V genes) (Fig. 3b-d and Supplementary Fig. 4) to be significantly associated with guQTLs in the HLA region. Some of these TCR genes were associated with the same lead guQTL, and we identified small networks of TRA and TRB genes being associated with the same set of *trans-*guQTLs (Fig. 3e). We also found pairs of TRA and TRB genes associated with the same lead guQTL (TRAV20, TRBV15 and rs601945; TRAV23/DV6, TRBV20-1 and rs9275184). The enrichment of genetic associations between HLA and TRA compared to TRB gene expression (mean 16.2 and 4.3 significant guQTLs per TRA and TRB gene, repectively) agrees with a previous *trans*-guQTL analysis^10^, and contrasts the seemingly bigger *cis*-acting effects compared to HLA-effects seen for many TRB genes^10,11^ and less extensive *cis*-effects for TRA compared to TRB genes observed in this study and reported by others in mixed T-cell populations^10^.

**Figure 3.**
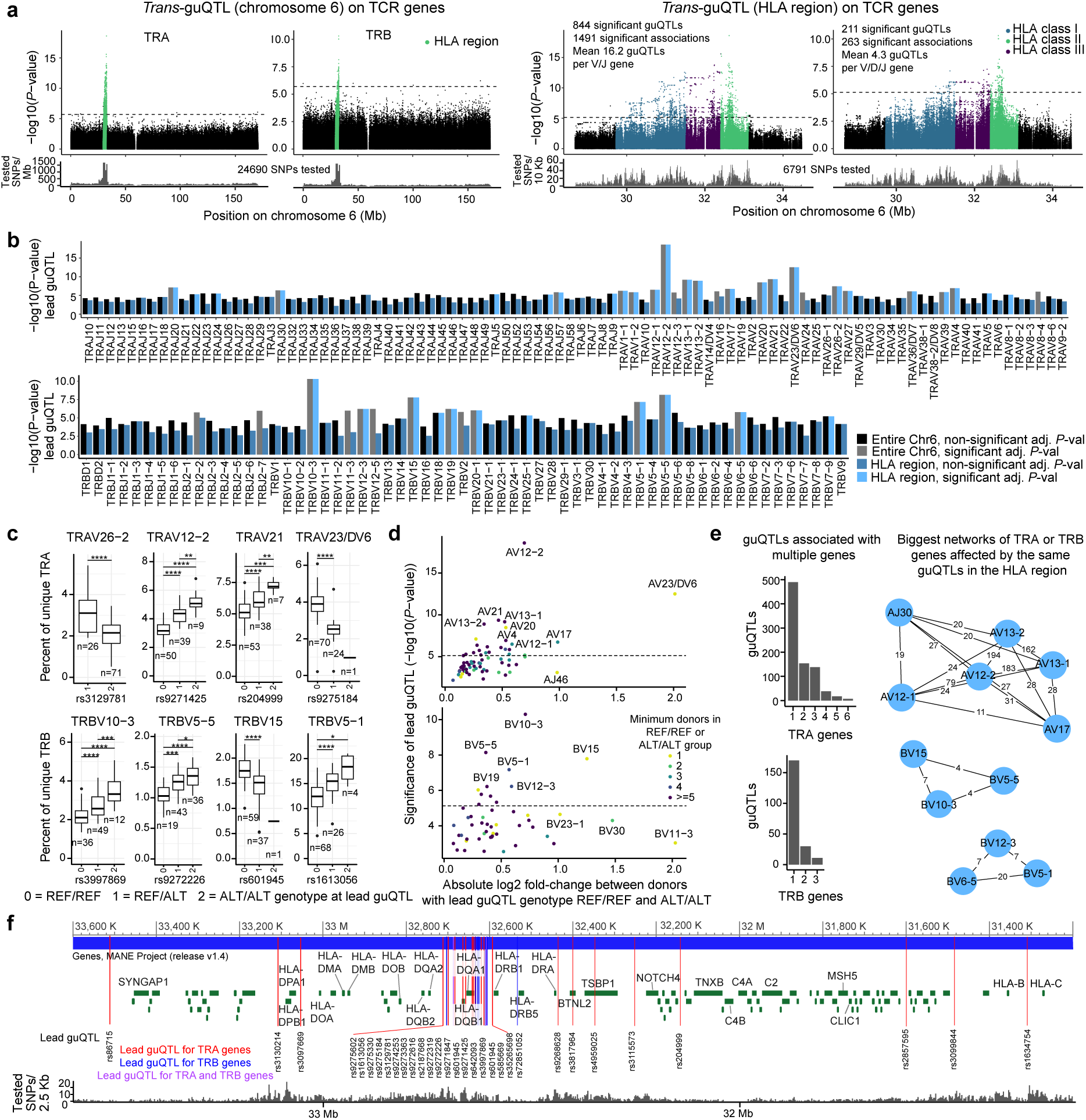
*Trans*-guQTL analysis of the influence of the HLA region on TRA and TRB gene usage in the naïve CD4^+^ TCR repertoire of control subjects. (**a**) Manhattan plots showing SNP associations for V, D and J genes with SNPs located on the entire chromosome 6 (left) or limited to 1 Mb before and after the HLA region (right). Dashed horizontal lines indicate significance threshold determined by Bonferroni correction. The lower tracks show number of tested SNPs along the chromosome. (**b**) Significance of the lead guQTL for each gene tested in (**a**). Black/grey bars correspond to the analysis including SNPs spanning the entire chromosome 6, blue bars correspond to the HLA-restricted analysis. (**c**) Examples of *trans-*mediated effects of lead guQTL genotype on TRA and TRB gene expression. Statistical significance was calculated using a Wilcoxon rank sum test followed by Bonferroni correction within each subfigure. Non-significant differences are not indicated. (**d**) Fold-change in expression between subjects with REF/REF and ALT/ALT genotype at lead *trans*-guQTL for each gene. (**e**) guQTLs associated with one or more TRA or TRB gene (left) and biggest networks of TRA genes and TRB genes associated to the same significant *trans*-guQTLs (right). (**f**) Location of significant lead *trans-*guQTLs within the HLA region of chromosome 6.

We also looked at where the significant lead guQTLs for TRA and TRB genes were located and observed that most were found within the HLA class II region, in particular between *HLA-DQA2* and *HLA-DRB1* (19/31 guQTLs) (Fig. 3f). This finding is in agreement with previous *trans*-guQTL analyses, placing most of the HLA class II effects in this region^10,12^.

### Gene usage frequency of many TCR genes varies depending on HLA-DQ allotypes

Based on this observation and motivated by the fact that HLA-DQ is implicated in CeD, we analyzed TCR gene usage in relation to HLA-DQ allotypes. The distribution of the allotypes in the control subjects is depicted in Fig. 4a. Restricting the analysis to the two most common allotypes DQ6 and DQ2.5, we found a striking association between HLA-DQ type and usage frequency for 12/20 and 7/16 of the most frequently used TRAV and TRBV genes, respectively. Examples included TRAV13*-*1 and TRBV5*-*1, in which we observed additive genetic effects on gene usage profiles associated with HLA genotypes (Fig. 4b,c and Supplementary Fig. 5). These findings indicate that variation within HLA influences selection of TCRs during thymic maturation of CD4^+^ T cells.

**Figure 4.**
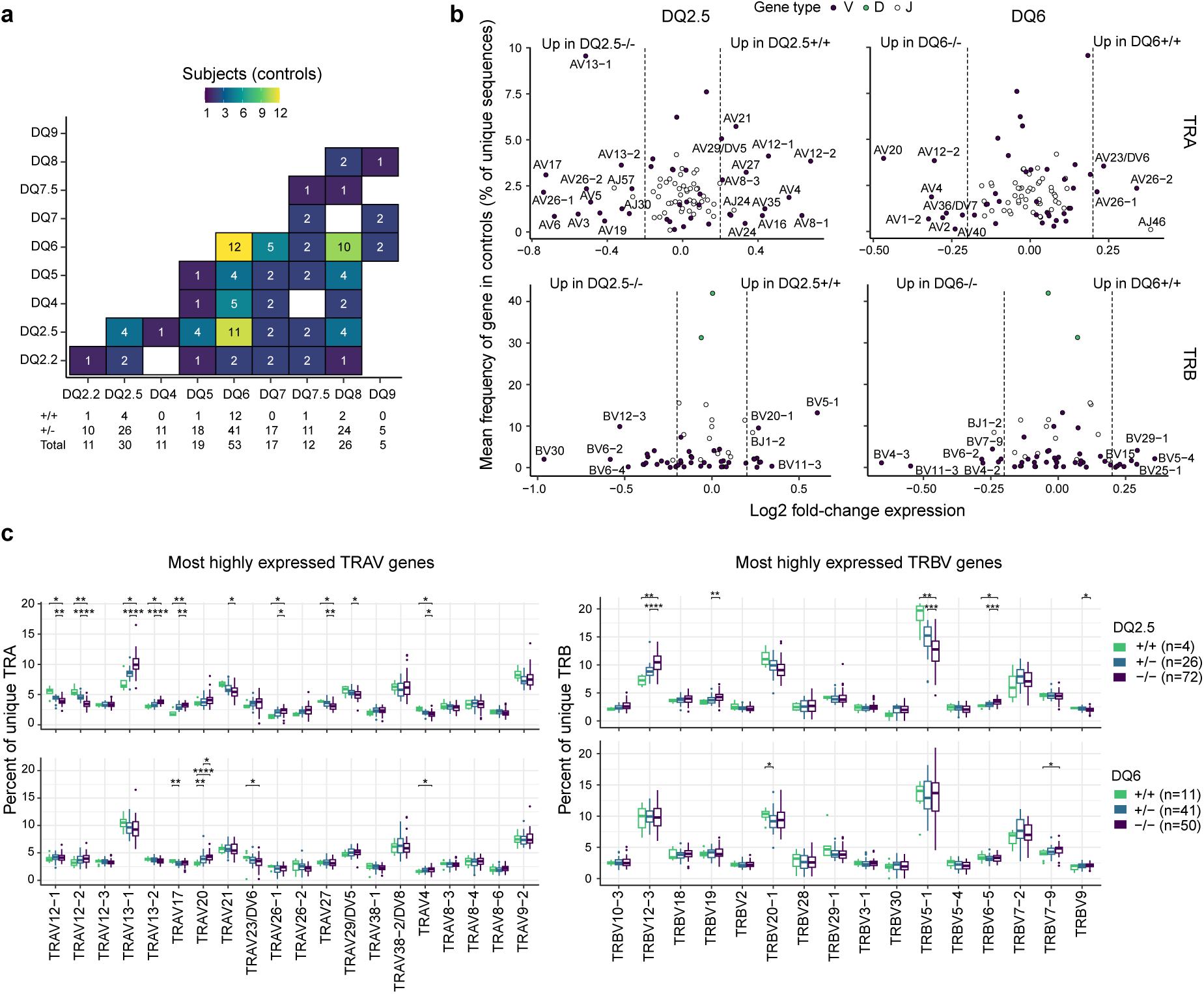
Variable TRA and TRB gene usage depending on HLA-DQ allotypes. (**a**) Distribution of HLA-DQ allotypes in control subjects, with heatmap indicating number of subjects with each combination of allotypes. The table below indicates total number of control subjects homozygous (+/+) or heterozygous (+/-) for each allotype, with total row indicating total number of donors with at least one copy of the allotype. (**b**) Log-fold change in expression of V (purple), D (green) and J (white) genes between subjects homozygous for or completely lacking the two most common HLA-DQ allotypes DQ2.5 (left) or DQ6 (right). Positive log-fold-change indicates more frequent usage in subjects homozygous for the allotype compared to subjects lacking the allotype. An absolute log2 fold-change of 0.2 is indicated with dashed vertical lines. (**c**) Distribution of gene usage frequency for most highly expressed TRAV (left) and TRBV (right) genes as a function of DQ2.5 (upper) or DQ6 (lower) allotypes. The number of donors in each allotype group are indicated to the right. Statistical significance was calculated using a Wilcoxon rank sum test followed by Bonferroni correction within each subfigure. Non-significant differences are not indicated.

### Individuals with HLA-DQ2.5 allotypes predisposing to CeD have increased frequency of CeD-relevant V genes in their naïve TCR repertoires

In CeD the T-cell response to gluten is stereotypical in that some selected genes are preferentially used for recognition of given epitopes and that this preferential usage is observed consistently across different individuals (Table 1)^9^. Interestingly, many of the V genes for which we observed increased frequency in the naïve TCR repertoires of HLA-DQ2.5^+^ control subjects are among those involved in these stereotyped responses of gluten-reactive T cells such as TRAV12-1, TRAV12-2, TRAV4 and TRBV5-1 (Fig. 5a,b and Table 1). This finding indicates that HLA-DQ2.5 may predispose to CeD not only by mediating presentation of deamidated gluten peptides in the gut, but also by increasing the frequency of T cells in the naïve repertoire that can respond to gluten by influencing the selection of T cells in thymus. The same strong effect was not seen for HLA-DQ8 for V genes commonly used by gluten-reactive effector T cells recognizing the immunodominant HLA-DQ8-glia-α1 epitope^29,30^ (Fig. 5c).

**Figure 5.**
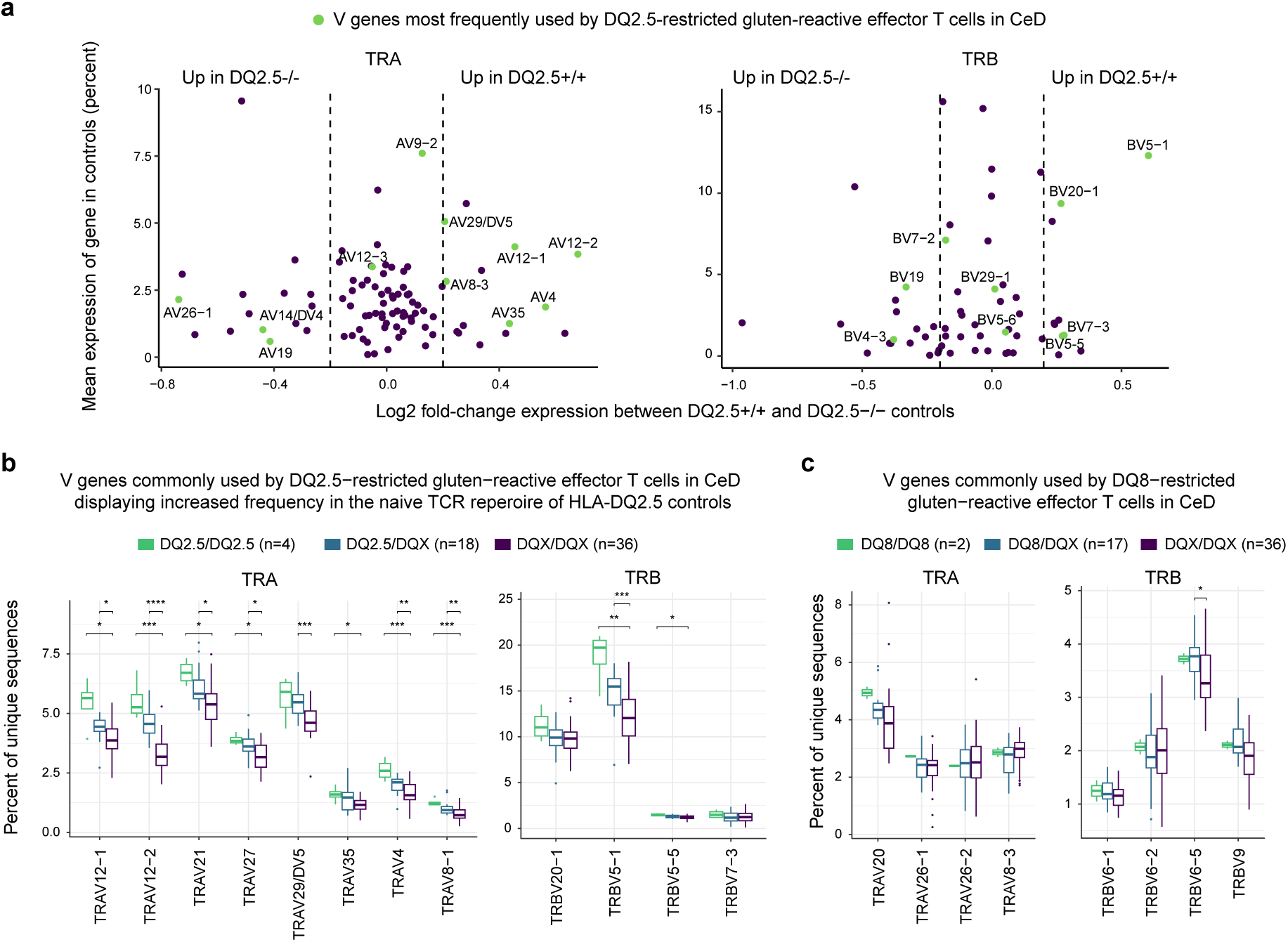
Subjects with the CeD-predisposing HLA-DQ allotype DQ2.5 have increased frequencies of several TRA and TRB gene segments commonly used by gluten-reactive T cells in CeD. (**a**) Log-fold change in expression of TRA (left) and TRB (right) V, D and J genes between control subjects homozygous for HLA-DQ2.5 (DQ2.5+/+) or lacking HLA-DQ2.5 (DQ2.5-/-). Positive log-fold-change indicates more frequent usage in DQ2.5+/+ compared to DQ2.5-/- subjects. An absolute log2 fold-change of 0.2 is indicated with dashed vertical lines. Genes most frequently used by DQ2.5-restricted gluten-reactive effector T cells in CeD^9^ are indicated in green. (**b**) Increased usage of many TRAV and some TRBV genes frequently involved in recognition of gluten epitopes by DQ2.5-restricted effector T cells in CeD^9^. Other CeD-associated HLA-DQ allotypes (DQ8, DQ2.2 and DQ7.5) were excluded from the analyses, meaning DQX indicates non-CeD-related allotypes. (**c**) Usage of some TRAV and TRBV genes frequently involved in recognition of gluten epitopes by DQ8-restricted effector T cells in CeD. Other CeD-associated HLA-DQ allotypes (DQ2.5, DQ2.2 and DQ7.5) were excluded from the analyses. Statistical significance was calculated using a Wilcoxon rank sum test for each gene and Bonferroni correction within each subfigure. Non-significant differences are not shown.

**Table 1.**
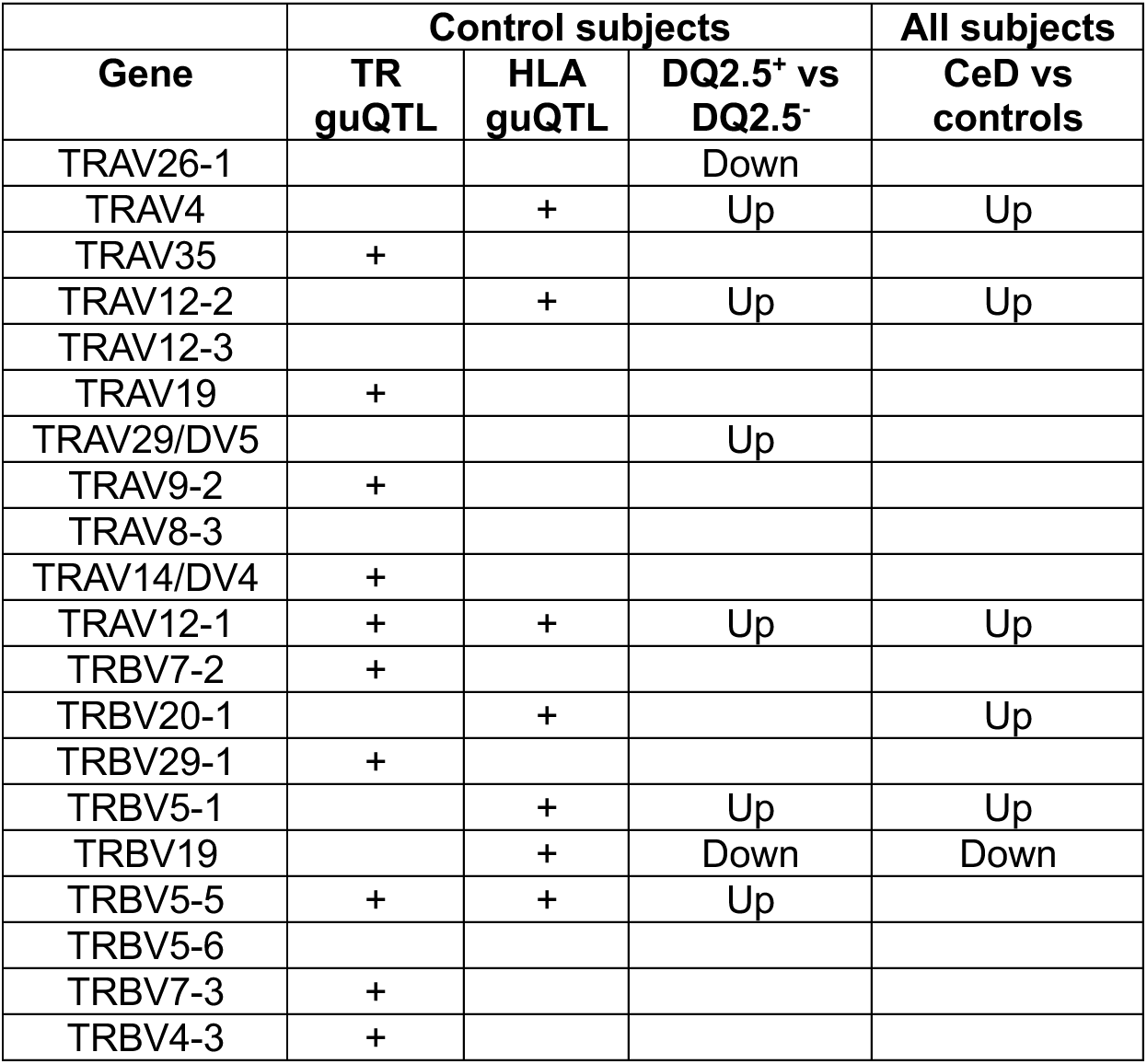
TRAV and TRBV genes most prominently involved in stereotypic recognition of gluten peptides bound to HLA-DQ2.5 in CeD.

These findings further our knowledge of how HLA variants, especially HLA-DQ2.5 in the case of CeD, predispose to disease. However, as HLA typically represents a necessary but not sufficient factor for disease development, it does not explain why most individuals with predisposing HLA allotypes do not develop a disease. One possibility could be that other genetic factors, such as variation within the TR loci, also predispose to disease by shaping the naïve TCR repertoires.

The genes listed are the top 11 TRAV and 9 TRBV genes used by HLA-DQ2.5-restricted gluten-reactive CD4^+^ effector T cells in CeD patients^9^. The columns indicate the results of the following analyses: TR guQTL - *cis*-guQTL analysis in controls (Fig. 2b). HLA guQTL - *trans*-guQTL analysis of HLA region in controls (Fig. 3b). DQ2.5^+^ vs DQ2.5^-^ - gene usage frequency partitioned by DQ2.5 allotype in controls (Fig. 4c). CeD vs controls - gene usage frequency in CeD subjects compared to controls (Fig. 6d). +: Significant effect on gene usage. Up: Significantly more frequently used in DQ2.5^+/+^ and/or DQ2.5^+/-^ compared to DQ2.5^-/-^ controls or in CeD compared to control subjects. Down: Significantly less frequently used in DQ2.5^+/+^ and/or DQ2.5^+/-^ compared to DQ2.5^-/-^ controls or in CeD compared to control subjects.

**Figure 6.**
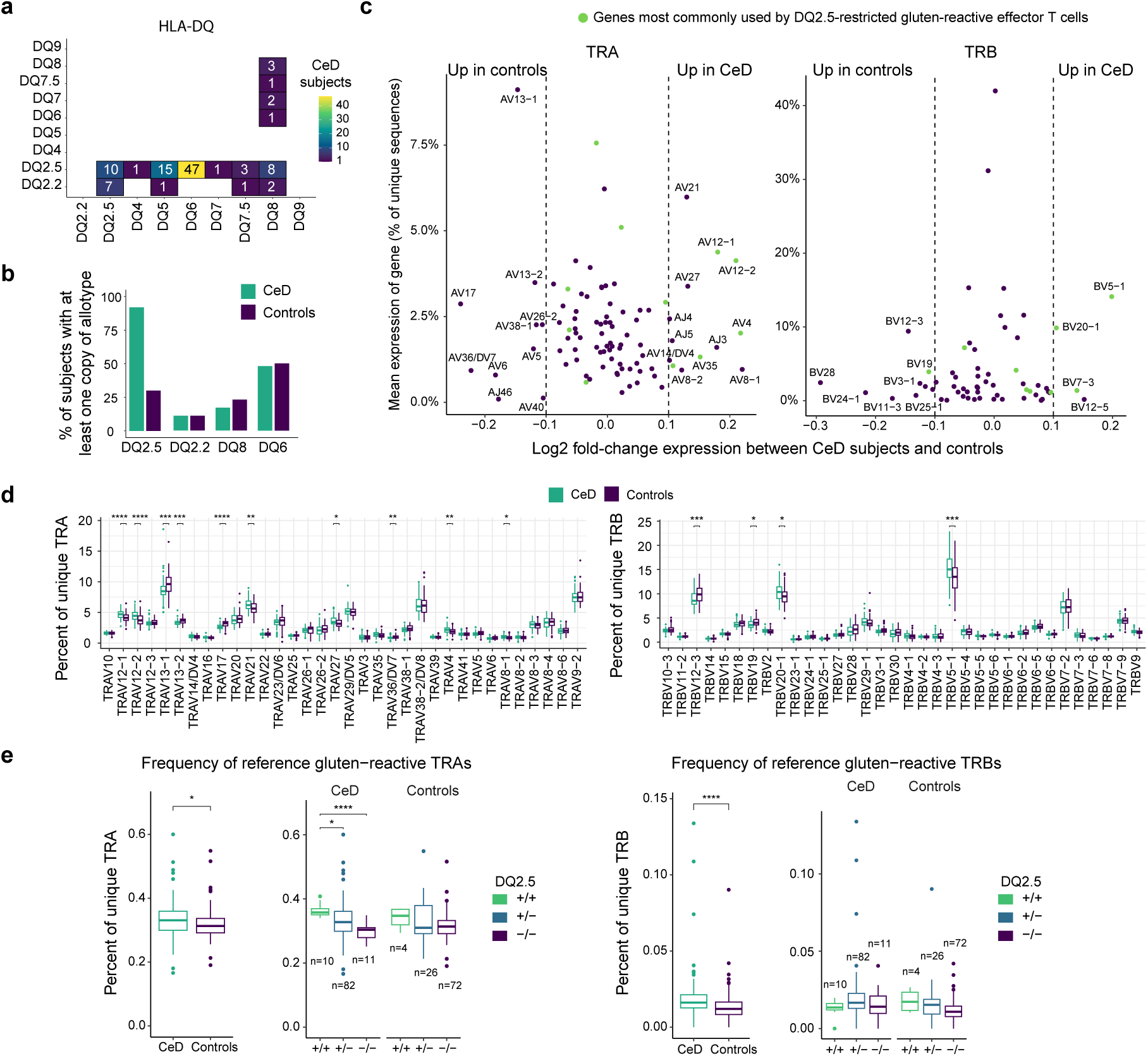
TRA and TRB gene usage frequencies in the naïve CD4^+^ TCR repertoire of CeD compared to control subjects. (**a**) Distribution of HLA-DQ allotypes in CeD subjects, with heatmap indicating number of subjects with each combination of allotypes. (**b**) Highly biased HLA-DQ allotype distribution in CeD compared to control individuals, with most CeD subjects expressing the DQ2.5 allotype (**c**) Log-fold change in expression of V, D and J genes between CeD and control subjects. Positive log-fold-change indicates more frequent usage in CeD subjects compared to controls. An absolute log2 fold-change of 0.1 is indicated with dashed vertical lines. Genes most commonly used by DQ2.5-restricted gluten-reactive effector T cells in CeD^9^ are highlighted in green. (**d**) Distribution of gene usage frequency for most highly expressed TRAV (left) and TRBV (right) genes in CeD subjects compared to controls. (**e**) Frequency of potentially gluten-reactive TRA (left) and TRB (right) sequences for CeD subjects compared to controls, based on a database of DQ2.5-restricted gluten-reactive TCR sequences from 50 CeD subjects^9^. Frequency is further conditioned by HLA-DQ2.5 allotype, with number of subjects in each group indicated on the graph. Statistical significance was calculated for each gene (**d**) or subfigure (**e**) using a Wilcoxon rank sum test and Bonferroni correction within each subfigure. Non-significant differences are not shown.

### Individuals with CeD have altered frequencies of TCR genes in their naïve CD4^+^ TCR repertoires compared to controls

Having included 103 subjects with CeD in this study, we set out to characterize potential differences between the naïve TCR repertoires of individuals with CeD and controls. Three of the CeD subjects were already diagnosed with CeD at the time of screening and were on a gluten-free diet, while the remaining 100 CeD subjects were diagnosed with CeD as a result of the screening procedure and follow-up appointments^20,21^.

Based on the observed impact of HLA class II polymorhisms on the composition of the naïve TCR repertoires of controls, we expected that the naïve TCR repertoires of CeD subjects and controls differ in terms of gene usage frequencies given the heavy skewing towards HLA-DQ2.5 in CeD individuals (Fig. 6a,b). Indeed, we observed large differences in gene usage frequency with higher expression of genes such as TRAV12-1, TRAV12-2, TRAV4 and TRBV5-1 in CeD subjects compared to controls (Fig. 6c,d and Supplementary Fig. 6). These genes were almost exclusively both CeD-associated and most frequently used in DQ2.5^+^ subjects (Table 1).

We further hypothesized that CeD subjects would have more gluten-reactive naïve T cells compared to controls due to their increased usage of many stereotypical V genes in CeD. We therefore identified potentially gluten-reactive naïve TCR sequences by matching to a reference database of TRA and TRB sequences derived from gluten-reactive HLA-DQ2.5-restricted CD4^+^ effector T cells from 50 CeD subjects^9^ (Supplementary Fig. 7). We found a total of 2061 unique potentially gluten-reactive sequences in the naïve CD4^+^ TCR repertoire of CeD individuals and controls (Fig. 6e). As expected, we observed a modest but significant increase of these sequences in CeD subjects compared to controls, which we considered was mainly a result of the V-gene bias associated with the DQ2.5 allotype. Specifically, we found increased frequency of potentially gluten-reactive TRAs, but not TRBs, in DQ2.5^+/+^ compared to DQ2.5^+/-^ or DQ2.5^-/-^ CeD subjects, potentially reflecting the relatively lower HLA-effect observed on TRB genes compared to TRA. Although not statistically significant, potentially gluten-reactive TRAs and TRBs were more frequent in DQ2.5^+/+^ compared to DQ2.5^+/-^ or DQ2.5^-/-^ controls.

### HLA explains many, but not all differences in naïve TCR repertoires between CeD subjects and controls

Our guQTL analyses revealed that the expression of most TCR genes were affected by either *cis-*effects or HLA-mediated *trans-*effects but also that several genes, such as TRAV26-2, displayed both types of effects (Fig. 7a,b). It is possible that more genes are affected by both *cis-* and *trans-*mediated effects, but that these were not strong enough to reach statistical significance or non-detectable due to SNP coverage.

**Figure 7.**
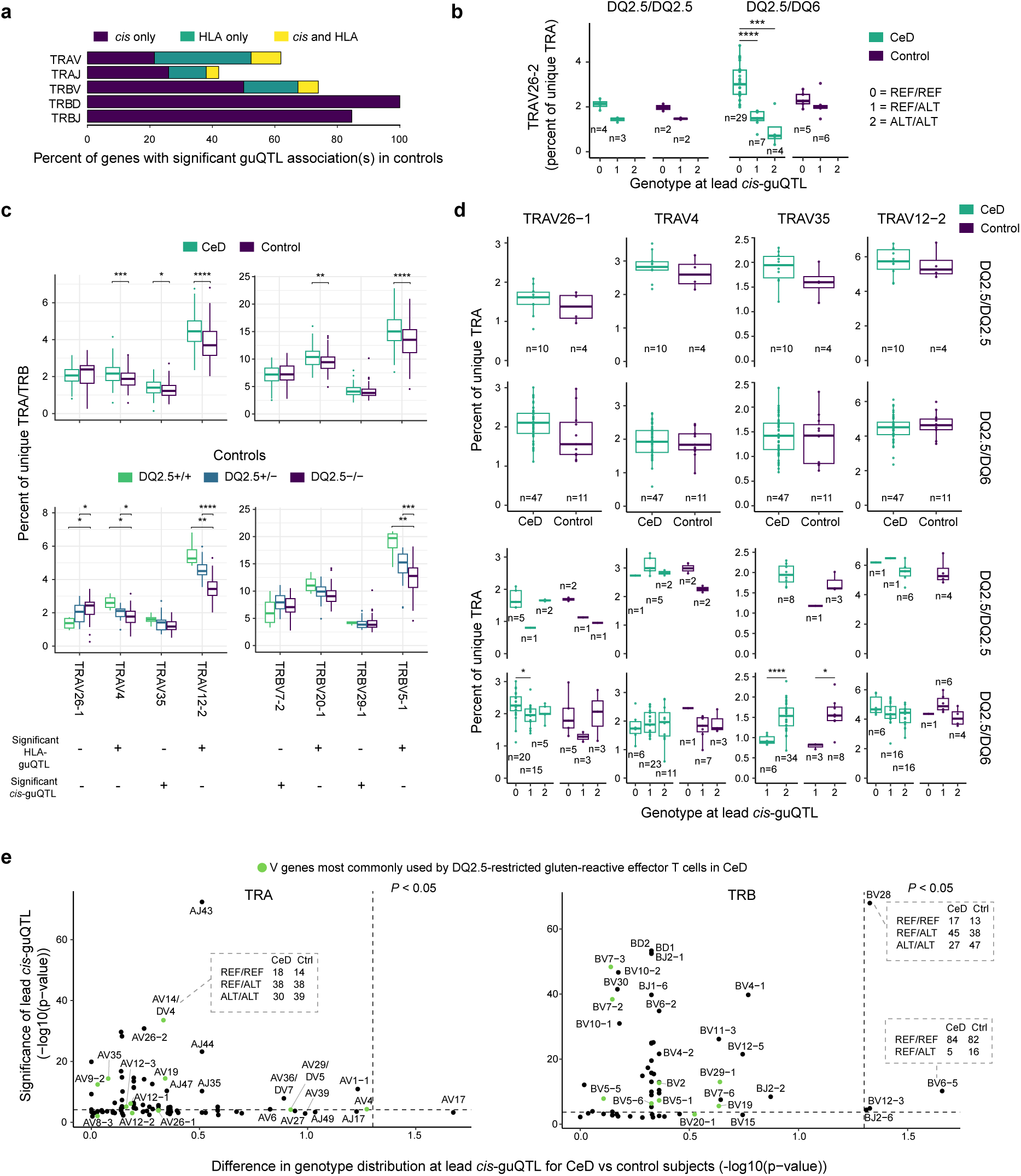
Relative contributions of *cis*- and HLA-mediated guQTL effects on usage of CeD-associated TCR genes. (**a**) Percentage of TRA and TRB genes with statistically significant *cis-* and/or *trans*(HLA)- mediated effects on usage of V, D or J genes. (**b**) TRAV26-2 usage conditioned on HLA-DQ type and lead *cis-*guQTL genotype. (**c**) Gene usage frequency for the four most CeD-relevant TRAV and TRBV genes. (**d**) The TRAV genes in (**c**) conditioned on HLA-DQ type and lead *cis-*guQTL genotype. Statistical significance was calculated with Wilcoxon rank sum tests (**b**-**d**). (**e**) Association analyses of lead *cis-* guQTL genotypes in CeD compared to control subjects. Horizontal dashed lines indicate significance threshold for *cis*-guQTL analysis after Bonferroni-correction. Vertical dashed lines indicate a significance threshold of *P =* 0.05 for the association of lead guQTL genotype in CeD subjects compared to controls. Examples of difference in lead guQTL genotype distributions are shown in dashed grey boxes.

Next, we set out to explain the differences in gene usage between CeD and control subjects (Fig. 6), in particular with regards to the V genes taking part in the stereotypic responses of gluten-specific T cells in CeD. These differences could most often be explained by HLA-related effects (Table 1 and Fig. 7c). To further investigate *cis-* effects, we controlled for HLA-mediated effects by performing conditional analyses limited to the HLA-DQ allotype combinations DQ2.5/DQ2.5 and DQ2.5/DQ6 (Fig. 7b,d and Supplementary Fig. 8). These analyses revealed that the *cis-*effects generally were the same in CeD and control subjects and persisted after reducing the confounding *trans*-mediated effects by HLA.

As some of the most CeD-relevant V genes had more pronounced *cis*-mediated rather than HLA-related effects on gene usage in the controls (Table 1), we hypothesized that CeD individuals and controls may have different usage of some of these genes due to the genotype at the lead *cis-*guQTLs. An association analysis identified one TRA and four TRB genes with significantly different lead guQTL genotype distributions in CeD individuals compared to controls (Fig. 7e). However, none of these genes were in the group of what was considered as particularly CeD-relevant V genes. Furthermore, several of the significantly associated guQTLs had weak if any effect on gene expression. An exception was the lead guQTL for TRBV28 which was significantly associated with disease status and had a highly significant *cis-*effect. The allele associated with low TRBV28 expression was more common in CeD individuals, explaining their seemingly lower expression of TRBV28 compared to controls (Fig. 6d). A limitation of the current study is the restricted number of CeD and control subjects studied. Thus, analysis of larger cohorts is needed for more definite answers as to whether a usage of CeD-relevant V genes can be coupled to *cis*-mediated effects.

## Discussion

Our results demonstrate that genetic variants both within the HLA and TR loci have significant effects on the naïve CD4^+^ TCR repertoire, and moreover that the HLA-DQ2.5 allotype which predisposes to CeD selects for usage of TCR genes being involved in stereotypical recognition of deamidated gluten epitopes by CD4^+^ T cells of CeD patients.

A strength of the study is that we carefully sorted naïve CD4^+^ T cells from peripheral blood, thus removing potentially confounding factors such as CD8^+^ T cells and/or antigen-experienced cells. In this manner we were able to study the output of thymic selection with regards to CD4^+^ T cells. Another strength is that our cohort consists of control subjects and CeD subjects, allowing us to investigate the effect of genetic variation in the HLA and TR loci both in health and in a disease setting with a well-characterized HLA-association and stereotypical TCR bias. By focusing on HLA in a *trans*-guQTL analysis, we provide compelling support in favor of the hypothesis that TCR-HLA compatibilities bias the usage of certain V genes during thymic selection^10^. It is conceivable that some V genes are less compatible with certain HLA variants, leading to more frequent apoptosis of the T cells carrying TCRs with these genes due to failed recognition of self-pHLA complexes during positive selection. It is also possible that TCR-HLA compatibilities may influence negative thymic selection if certain TCR genes are more likely to form TCRs that bind self-pHLA too strongly. The observation that HLA-mediated *trans*-effects were almost exclusively seen for V genes further underlines the importance of germline-encoded residues in the V genes for HLA contact and selection. Our *trans*-guQTL analysis revealed stronger HLA-effects for TRA compared to TRB genes, in line with previous data from unsorted peripheral blood T cells^10^. This observation speaks to involvement of TRA in interaction with HLA at least when looking at germline-encoded residues.

A key finding is that controls carrying the CeD-predisposing allotype HLA-DQ2.5 have increased frequency of several TRAV and TRBV genes in their naïve CD4^+^ TCR repertoires that are employed in the stereotypic recognition of gluten epitopes by disease-driving CD4^+^ T cells in CeD. This suggests that HLA has a dual predisposing role in CeD in (1) antigen presentation by allowing efficient presentation of deamidated gluten peptides and (2) skewing the naïve TCR repertoire towards T cells that are capable of recognizing gluten peptides presented on HLA-DQ2.5. Previous literature has described the structural basis for selection of certain V genes such as TRAV4, TRBV20-1 and TRBV29-1 in the gluten-reactive effector T cells in CeD, with interactions between germline-encoded residues in the TCR and HLA-DQ2.5^31^. Given the increased usage we observed in HLA-DQ2.5^+^ individuals in the naïve TCR repertoire for at least some of these genes, it is likely that these V genes are selected in the same manner during thymic selection. Due to the generally low frequency of any antigen-specific TCR in the naïve repertoire, any increase in this frequency may predispose to disease development by increasing the chance of the antigen-specific cell seeing their antigen. It is conceivable that HLA may predispose to other diseases in a similar manner by skewing the naïve TCR repertoires.

One limitation of our study is that it lacks single-cell resolution and thus information about chain pairing. TRB and TRA are not selected completely independently – TRA needs to be able to pair with TRB, and it is the combined affinity to pHLA that will determine the outcome of selection. Lack of chain pairing information may also affect our ability to detect truly gluten-reactive TCRs based on reference sequences. Nevertheless, increased detected frequencies of both CeD-relevant TRA and TRB chains likely increases the chances of forming gluten-reactive TCRs. Our detection of gluten-reactive TCR sequences is further limited by our reference dataset not being exhaustive.

In addition to identifying HLA-mediated *trans*-effects on TCR gene usage, we also identified extensive *cis*-effects, especially for the TRB locus. The apparent higher degree of *cis*-effects on TRB gene usage compared to TRA agrees with previous *cis*-guQTL analyses based on unsorted PBMCs^10^. Our results further verify previous reports of specific *cis*-guQTLs within the TR loci^10^, such as the strong *cis*-effects on TRAJ24 and TRAJ28 known from studies of narcolepsy^17^. In contrast to the HLA-mediated *trans*-effects, both V, D and J genes were commonly affected by *cis*-guQTLs.

Notably, we found that most of the guQTLs were located outside the genes themselves, but commonly close by. While it is possible that some of these guQTLs serve as markers for genetic variants in exons, it is likely that many of the *cis*-acting genetic effects on TRA and TRB gene usage come from regulatory elements within the TR loci. Due to the relatively small number of tested SNPs across the TRA and TRB loci it is hard to pinpoint exactly where the causative genetic variants are located.

Taken together, we show that the naïve CD4^+^ TCR repertoire is influenced both by genetic variants acting in *cis* and in *trans*. Based on the knowledge of which TCR genes are involved in recognition of gluten peptides by disease-driving T cells in CeD, we are able to make the case that HLA by acting in *trans* predisposes to a human disease by selecting the naïve TCR repertoire.

## Acknowledgements

We would like to thank all participants of the HUNT4 study for donation of blood samples to this project, and the personnel at HUNT Research Centre and Levanger Hospital for assistance with collecting the sample material of the study subjects. The Trøndelag Health Study (HUNT) is a collaboration between HUNT Research Centre (Faculty of Medicine and Health Sciences, Norwegian University of Science and Technology [NTNU]), Trøndelag County Council, Central Norway Regional Health Authority and the Norwegian Institute of Public Health. We thank Marie Kongshaug Johannessen, Marte Viken and Lisa Brynjulfsen for assistance with experimental procedures. The Norwegian Sequencing Centre, the Flow Cytometry Core Facility and the Section for Transplantation Immunology, Department of Immunology (both at Oslo University Hospital) provided analytical service. Sigma2 provided computational resources. Data analysis was performed on the TSD (Tjenester for Sensitive Data) facilities, owned by the University of Oslo. Figure 1a was made with Biorender. This study was funded by means from the South-Eastern Norway Regional Health Authority (project 2022040) and the University Oslo (to L.M.S), from the Norwegian Coeliac Society (to L.M.S. and E.N.-J) and from NTNU’s Outstanding Academic Fellows Program (to E.N.-J.).

## Author contributions

L.M.S. conceived and designed the study. I.L. and S.D.-K. contributed to the study design. I.L., A.O. and S.D.-K. performed experiments. I.L. performed the formal data analysis and created the figures. I.L., L.M.S. and C.T.W. interpreted the data and provided input to data analysis. R.H. provided HLA-DQ typing results. E.N.-J. and K.E.A. provided biological samples and ethical approvals. S.D.-K. and L.F.R. contributed to sample collection. L.M.S. and E.N.-J. provided funding. L.M.S. supervised the study. I.L. drafted the original manuscript. I.L., L.M.S. and C.T.W. revised the manuscript. All authors approved the revised manuscript.

## Methods

### Ethics statement

This study was approved by the Regional Ethics Committee of South-Eastern Norway (REK 6544). All subjects gave their informed, written consent before sample collection. Cryopreserved peripheral blood mononuclear cells (PBMCs) were stored in the biobank “Tarmsykdommer” at Oslo University Hospital (REK 20521).

### Study participants

We obtained cryopreserved PBMCs from 103 subjects diagnosed with CeD and 103 age-matched controls from subjects who participated in the fourth population-based Trøndelag Health Study (HUNT4)^20–22,32^. Except for three CeD subjects with preexisting CeD diagnosis and following a gluten-free diet, all other CeD subjects were newly diagnosed CeD cases identified by serological screening of the HUNT4 population and follow-up clinical assessments confirming the diagnosis based on duodenal biopsies.

### Magnetic bead separation and fluorescence-activated cell sorting (FACS)

Cryopreserved PBMCs were thawed in RPMI-1640 supplemented with 20% fetal calf serum (FCS) and washed with phosphate buffered saline (PBS) supplemented with 1% FCS and 2 mM EDTA. Depending on cell numbers, samples were subjected to magnetic bead separation followed by FACS (strategy 1), or only FACS for samples with low cell numbers (strategy 2) (Supplementary Fig. 1a). For magnetic bead separation with MS Columns (Miltenyi Biotec), CD19^-^ CD4^+^ cells (as well as CD19^+^ cells included for a separate study) were enriched for using human CD19 MicroBeads (Miltenyi Biotec) followed by human CD4 MicroBeads (Miltenyi Biotec) for the CD19-depleted fraction, and CD19-enriched cells were pooled together with CD4-enriched CD19-depleted cells for FACS. The CD19-depleted CD4-depleted fraction was frozen down at −20°C as cell pellets for extraction of genomic DNA (gDNA). For samples sorted directly without bead separation, CD19^-^ CD4^-^ cells for gDNA extraction were instead obtained by FACS. When insufficient numbers of naïve T cells could be sorted from the starting material, additional PBMCs from the same subject were thawed when available and used directly for an additional round of sorting (strategy 3). These cells had already been thawed once and depleted for CD4^+^ effector memory T cells using the human CD4+ Effector Memory T Cell Isolation Kit (Miltenyi). Prior to FACS cells were resuspended in flow buffer (PBS supplemented with 2% FCS and 1 mM EDTA), and stained on ice with LIVE/DEAD Fixable Aqua Dead Cell Stain Kit (Invitrogen) and the appropriate antibody mix according to sorting strategy (Supplementary Table 1). Cells were washed with flow buffer and bulk sorted on a FACSMelody (BD) cell sorter at the Flow Cytometry Core Facility at Oslo University Hospital (location Gaustad or Ullevål) and collected in RNase/DNase-free microcentrifuge tubes containing 100 μl PBS supplemented with 50% FCS. Sorted naïve T cells (live CD4^+^ CD62L^+^ CD45RA^+^ CD19^-^ CD14^-^) were washed in PBS, and pellets were snap-frozen in liquid nitrogen and stored at −72°C until RNA extraction. Washed cell pellets for gDNA extraction were stored at −20°C. Purity of all samples was checked after sorting, and flow cytometry data were analyzed with FlowJo v10.8.

### Isolation of RNA

Total RNA was extracted from naïve T cells using the RNeasy Micro Kit (Qiagen) for samples up to 0.5 million cells or RNeasy Mini Kit (Qiagen) followed by upconcentration using the RNeasy MinElute Cleanup Kit (Qiagen) for larger samples. RNA was eluted in 14 μl RNase-free H_2_O, and the purity and concentration of the isolated RNA was measured by Nanodrop One or Nanodrop 1000 (Thermo Fisher). The integrity and concentration of 22 of the samples were additionally measured on an Agilent 2100 Bioanalyzer instrument with the Agilent RNA 6000 Pico kit to ensure an overall good sample quality (Supplementary Fig. 1d). Isolated RNA was stored at −72°C until library preparation.

### Isolation, purification and quantification of gDNA

Cells for gDNA extraction were obtained by magnetic bead depletion of CD19^+^ and CD4^+^ cells and/or by sorting of CD19^-^ CD4^-^ cells by FACS. gDNA was extracted from frozen cell pellets using the QIAamp DNA Mini Kit (Qiagen). The gDNA concentration and purity was initially measured by Nanodrop. Low concentration gDNA samples or samples with low purity (A_260_/A_230_ < 1.5 or A_260_/A_280_ < 1.8) were further upconcentrated and purified as suggested by the Axiom 2.0 gDNA Sample Preparation Guideline, using 0.5 volumes of 7.5 M NH_4_OAc (Sigma-Aldrich) and 2.5 volumes of absolute ethanol. The integrity of gDNA samples was confirmed by running approximately 100 ng of each sample on a 1% agarose gel next to 6 μl of GeneRuler 1kB DNA Ladder (Thermo Fisher) (Supplementary Fig. 2a). For a small number of subjects with insufficient numbers of isolated CD19^-^ CD4^-^ cells or degraded, low-concentration or poor quality gDNA, unsorted PBMCs were thawed and used directly for gDNA extraction.

### SNP genotyping

For 196 of the subjects, the gDNA was used for SNP genotyping with an Axiom Human Genotyping SARS-CoV-2 Research Array (Thermo Fisher). gDNA samples were subsequently quantified in duplicate reactions using the PicoGreen dsDNA Quantification Kit (Molecular Probes) according to the manufacturer’s instructions, and normalized to 5 ng/μl. The Axiom arrays were prepared and read at the Centre for Integrative Genetics (Cigene) at the Norwegian University of Life Sciences, and the resulting CEL files were processed with the Axiom Analysis Suite using the “Best Practices Workflow” pipeline (Supplementary Fig. 2b). One sample could not be read during data acquisition, and an additional eight samples together with all negative controls failed quality control and were discarded. Further quality control was performed by matching known compared to calculated sex (Supplementary Fig. 2c), manual inspection of a selection of SNPs, analysis of intentionally included sample replicates and the degree of uniqueness between unrelated samples (Supplementary Fig. 2d), and inspection of potential batch effects.

### HLA inference and HLA-DQ typing

From the “best and recommended” SNPs, SNPs in the “HLA inference” module were extracted and used for HLA inference with Axiom HLA Analysis v1.2 (Thermo Fisher). HLA-DQ allotypes were manually assigned based on the inferred *HLA-DQA1* and *HLA-DQB1* alleles and when in doubt applying additional information derived from HLA-DRB1 inference and the known linkage disequilibria between HLA-DR and HLA-DQ allotypes. Additionally, four healthy controls and all but one CeD subject from the HUNT Study had previously undergone HLA-DQ typing by sequencing exons 2 and 3 of the *HLA-DQA1* and *HLA-DQB1* genes at HistoGenetics (Ossining, NY, USA). High-resolution typing was performed using paired-end Illumina Next Generation Sequencing, with further phasing and rare variant resolution achieved via PacBio platforms. Quality control was conducted through HistoGenetics’ standard analytical pipeline. Relevant HLA-DQ allotypes were inferred from results reported as G or MAC/NMDP codes. The robustness of the HLA inference was verified by comparing the inferred to the sequenced HLA-DQ types when both methods had been used. Five controls with no or unreliable HLA-DQ typing and/or inference were additionally HLA-DQ typed with a One Lambda LABType SSO Class II DQA1/DQB1 Typing test (Thermo Fisher) at the Department of Immunology and Transfusion Medicine at Oslo University Hospital.

### Preparation of naïve TCR Illumina sequencing libraries (AIRR-seq)

RNA samples from naïve CD4^+^ T cells were used for preparation of in total seven Illumina amplicon sequencing libraries organized by descending RNA input so that libraries with the lowest RNA input quantity contained the most samples. Each library consisted of equal numbers of controls and CeD subjects, with CeD samples interspersed with controls during library preparation to avoid batch effects. Library preparation was performed with a 5’ RACE strategy and semi-nested PCR approach, using a modified version of a previously described protocol^33^. All primer sequences are listed in Supplementary Table 2. RNA was mixed with 2 mM dNTP (Thermo Fisher), 2 μM oligo d(T) and 2 U/μl RNase inhibitor (New England Biolabs) in 8-well PCR strips and incubated at 72°C for three minutes before cooling down on ice. A reverse transcription (RT) mix was added to the samples, resulting in a final concentration of 1 mM dNTP, 1μM oligo d(T), 1.5 U/μl RNase inhibitor, 0.8 M betaine (Sigma-Aldrich), 6 mM MgCl_2_ (Sigma-Aldrich), 1x FS buffer (Invitrogen), 2.5 mM DTT (Invitrogen), 5 U/μl SuperScriptII (Invitrogen) and 2 μM template switch oligo (TSO) containing 12 nt unique molecular identifiers (UMIs). RT was performed under the following cycling conditions: 90 min × 42°C, 15 min × 72°C, 4°C hold.

TRA and TRB transcripts were amplified in three subsequent rounds of PCR, with each reaction performed in triplicates in 96-well plates. In the first round (PCR1), the synthesized cDNA was split into triplicate reactions for each sample, with a final concentration of 1x KAPA HiFi HotStart ReadyMix (Kapa Biosystems), 40 nM STRT-fwd-Long oligo, 200 nM STRT-fwd-Short oligo, 200 nM TRAC oligo and 200 nM TRBC oligo. PCR1 was performed with the following conditions: 1 min × 98°C, 5 × (10s × 98°C, 60 s × 72°C), 5 × (10s × 98°C, 30 s × 70°C, 40 s × 72°C), 8 × (10s × 98°C, 30 s × 68°C, 40 s × 72°C), 4 min × 72°C, 4°C hold. For samples with the lowest RNA input (last two libraries), the number of cycles in the last amplification round was increased from 8 to 12 to ensure sufficient amplification.

Each 1^st^ PCR reaction was used as template in two separate PCR2 reactions, one for TRA and one for TRB, using 1 μl of template in each reaction with a final concentration of 1x KAPA HiFi HotStart ReadyMix, 200 nM R2_PCR2_InX oligo, and 200 nM R1_PCR2_TRACX or R1_PCR2_TRBCX oligo, where X indicates sample-specific barcodes (Supplementary Table 3). A dual indexing strategy was used. Index sequences were derived from Islam et al.^34^, and each 6 nt index differed by at least three nucleotides. PCR2 was performed with the following conditions: 1 min × 98°C, 10 × (10s × 98°C, 20 s × 65°C, 30 s × 72°C), 4 min × 72°C, 4°C hold. 2^nd^ PCR products were further amplified in PCR3, using 2 μl of template in each reaction with a final concentration of 1x KAPA HiFi HotStart ReadyMix, 200 nM R1 oligo and 200 nM R2 oligo to introduce Illumina sequencing adapters. PCR3 was performed with the following conditions: 2 min × 95°C, 15 × (20s × 98°C, 30 s × 60°C, 40 s × 72°C), 5 min 72°C, 4°C hold.

Triplicate 3^rd^ PCR products were pooled together and visualized on a 1% agarose gel to confirm the presence of amplicons of correct size for each sample. Equal volumes of each sample in a library were subsequently pooled, keeping TRA and TRB reactions separate, purified with 0.65x AMPure XP beads (Beckman Coulter) and eluted in Monarch DNA Elution Buffer (New England Biolabs). The concentration and purity of pooled TRA and TRB libraries were measured using Nanodrop, and successful removal of small DNA bands was visualized on a 1% agarose gel. Equal concentrations of TRA and TRB from a given library were pooled for sequencing on an Illumina NovaSeq 6000 instrument at the Norwegian Sequencing Centre using paired-end reads of length 250 bp.

### Processing of raw AIRR-seq data

Raw paired-end fastq sequences (“R1” and “R2”) were processed with the pRESTO^35^ tool suite on the TSD (Tjenester for Sensitive Data) facilities, a platform for secure storage and processing of sensitive data. Reads were first demultiplexed according to sample indices detected with the *MaskPrimers score* function, allowing one nucleotide mismatch. For each subject in a sequencing library, demultiplexed reads with the correct R1 and R2 indices were trimmed to *Q* = 30 with *FilterSeq trimqual*, and reads < 150 bp (R1) or 75 bp (R2) were discarded. Reads containing the expected primer sequences with an error rate of 0.2 were retained, and these primer sequences were cut. The 12 base UMIs were extracted from R2 reads and assigned to R1 by pairing R1 and R2 reads, and unpaired reads were discarded. R1 or R2 reads with the same UMI were grouped and aligned with *ClusterSets* using vsearch v.2.3.2 and *AlignSets* using muscle v3.8.31, and consensus sequences were generated with *BuildConsensus* for each UMI set. Only R1 consensus sequences which could be successfully paired to an R2 consensus sequence were retained, and sequences with only one supporting read were discarded. The resulting R1 sequences (containing the 3’ end of the V gene, CDR3, the J gene and parts of TRAC or TRBC) were annotated with the Change-O tool suite^36^ using IgBlast^37^ v.1.19.0 and a reference database of TR sequences downloaded from the international ImMunoGeneTics information system (IMGT)^38,39^ in October 2022. Sequences recognized by IgBlast as rearranged TRA or TRB sequences were further trimmed to only contain the V(D)J sequence. Identical V(D)J sequences were collapsed, and only sequences with at least two unique UMIs were retained. To greatly reduce the impact of technical noise, such as PCR or sequencing errors or incomplete transcription leading to different transcript lengths, the *DefineClones* function of Change-O was used to identify sequences with overlapping V- and J-gene assignments and identical CDR3 nucleotide sequences. Non-productively rearranged V(D)J sequences were discarded from the analyses. Additionally, sequences covering fewer than 70 nucleotides of the V gene were discarded to reduce uncertain gene annotations.

### Quantification of TCR gene usage

The gene usage frequency of each V, D, or J gene was calculated separately for TRA and TRB sequences with the *countGenes* function (in *gene* mode) of Alakazam^36^ in R v4.4.3 or v4.5.0. In order to only consider truly uniquely rearranged sequences, sequences belonging to the same “clone” (likely arising from technical noise, see above) were only counted once. Only TRA or TRB genes were considered. Genes detected in fewer than three subjects were discarded from the analysis.

### Quantification of potentially gluten-reactive TRA and TRB sequences

A reference database of 2562 unique gluten-reactive TRA and 2428 unique TRB sequences was obtained from single-cell AIRR-seq of HLA-DQ2.5-restricted gluten-reactive effector memory T cells isolated from the intestinal lamina propria and/or peripheral blood from a total of 50 CeD subjects^9^ (Supplementary Fig. 7). Any naïve TRA or TRB with identical CDR3 amino acid sequence to a reference sequence and additionally shared overlapping V- and J-gene assignments, was considered a potentially gluten-reactive sequence. The total frequency of potentially gluten-reactive naïve TRA and TRB sequences in each subject was calculated with the with the *countGenes* function of Alakazam, again only counting sequences belonging to the same “clone” once.

### Gene usage QTL analysis

For all guQTL analyses only SNPs defined as “best and recommended” by the Axiom *Best Practices Workflow* and passing the following criteria were retained: Minor allele frequency > 0.05, non-hemizygous, Hardy-Weinberg equilibrium *P* > 0.000001. For *cis*-guQTL SNPs located within 1 million bases (Mb) of the beginning and end of the TRA locus on chromosome 14 or TRB locus on chromosome 7 according to the GRCh38.p14 (hg38) assembly were kept. For *trans*-guQTL analyses of HLA effects on TRA or TRB gene expression, SNPs on chromosome 6 (first approach) or within 1 Mb of the beginning and end of the HLA region on chromosome 6 (second approach) according to GRCh38.p14 were kept. guQTL analyses of V, (D) and J gene expression were conducted separately for TRA, TRB and the HLA region. Association tests were performed using linear regression, and we corrected for multiple hypothesis testing with Bonferroni correction on a per-gene level.

### Network analysis of shared guQTLs

Networks of multiple guQTLs significantly associated with a group of TRA or TRB genes were drawn using networkx in python v3.8, with genes represented as nodes and shared guQTLs represented as edge weight. The networks were pruned to edge weights > 2 (TRA), > 10 (TRB) or > 3 (HLA region), and subgraphs of genes with high degree of shared guQTLs were identified by manual inspection and drawn separately for improved visualization.

### Visualization of gene and SNP locations

TRA, TRB and HLA genes and their position in relation to tested SNPs were visualized using the National Center for Biotechnology Information (NCBI)^40^ Genome Data Viewer, using the GRCh38.p14 (hg38) reference assembly and NCBI RefSeq Annotation GCF_000001405.40-RS_2024_08. Gene tracks were cleaned up in Adobe Illustrator to improve readability and remove non-functional TRA and TRB genes for simplicity.

### Statistical analyses

All statistical tests throughout this paper were two-tailed. Statistically significant differences in gene expression between two sample groups were identified using the Wilcoxon rank sum test followed by Bonferroni correction within each subfigure. Differences between CeD and control subjects related to subject characteristics and AIRR-seq library construction were also tested with the Wilcoxon rank sum test. Throughout the article, *P* values are reported with the following significance levels: * = *P* < 0.05, ** = *P* < 0.01, *** = *P* < 0.001, **** = *P* < 0.0001, ns = *P* ≥ 0.05. Boxplots are drawn with ggplot2 and show either all data points or only outliers as well as the following statistics: Median, hinges corresponding to the first and third quartiles, and whiskers indicating the highest or lowest value no further than 1.5 × the inter-quartile range from each hinge. Genotype distributions at each lead guQTL were tested for association to subject disease status (CeD or control) by Chi-squared tests (lead guQTLs with at least five subjects in each genotype group for each disease status) and Fisher’s exact tests (lead guQTLs with fewer than five subjects in at least one of the genotype groups).

## Data availability statement

Raw and processed AIRR-seq data, as well as genotyping data for the TRA, TRB and HLA regions and necessary metadata will be deposited at the European Genome-Phenome Archive (EGA) under controlled access due to data sensitivity. Data access is regulated through a data access agreement, and usage of the data will be limited to studies seeking to advance the understanding of immunological mechanisms for autoimmune diseases. Inquiries related to data access may be directed to the corresponding authors. A formal data access requests must be made throught the EGA website, and will be evaluated by a Data Access Committee (consisting of L.M.S., I.L. and E.N.-J.) within two weeks. Data analysis scripts are available from the authors upon reasonable request.

## References

1. Vyse, T.J. & Todd, J.A. Genetic analysis of autoimmune disease. Cell 85, 311–8 (1996).

2. Sollid, L.M., Pos, W. & Wucherpfennig, K.W. Molecular mechanisms for contribution of MHC molecules to autoimmune diseases. Curr. Opin. Immunol. 31, 24–30 (2014).

3. Parkes, M., Cortes, A., van Heel, D.A. & Brown, M.A. Genetic insights into common pathways and complex relationships among immune-mediated diseases. Nat. Rev. Genet. 14, 661–73 (2013).

4. Dendrou, C.A., Petersen, J., Rossjohn, J. & Fugger, L. HLA variation and disease. Nat. Rev. Immunol. 18, 325–339 (2018).

5. Thorsby, E. Invited anniversary review: HLA associated diseases. Hum. Immunol. 53, 1–11 (1997).

6. Jabri, B. & Sollid, L.M. T cells in celiac disease. J. Immunol. 198, 3005–3014 (2017).

7. Kim, C.Y., Quarsten, H., Bergseng, E., Khosla, C. & Sollid, L.M. Structural basis for HLA-DQ2-mediated presentation of gluten epitopes in celiac disease. Proc. Natl. Acad. Sci. USA 101, 4175–9 (2004).

8. Molberg, O. et al. Tissue transglutaminase selectively modifies gliadin peptides that are recognized by gut-derived T cells in celiac disease. Nat. Med. 4, 713–7 (1998).

9. Dahal-Koirala, S. et al. Comprehensive analysis of CDR3 sequences in gluten-specific T-cell receptors reveals a dominant R-motif and several new minor motifs. Front. Immunol. 12, 639672 (2021).

10. Sharon, E. et al. Genetic variation in MHC proteins is associated with T cell receptor expression biases. Nat. Genet. 48, 995–1002 (2016).

11. Russell, M.L. et al. Combining genotypes and T cell receptor distributions to infer genetic loci determining V(D)J recombination probabilities. eLife 11, e73475 (2022).

12. Gao, K. et al. Germline-encoded TCR-MHC contacts promote TCR V gene bias in umbilical cord blood T cell repertoire. Front. Immunol. 10, 2064 (2019).

13. Klarenbeek, P.L. et al. Somatic variation of T-cell receptor genes strongly associate with HLA class restriction. PLoS One 10, e0140815 (2015).

14. Ishigaki, K. et al. HLA autoimmune risk alleles restrict the hypervariable region of T cell receptors. Nat. Genet. 54, 393–402 (2022).

15. Zahid, H.J. et al. Large-scale statistical mapping of T-cell receptor β sequences to Human Leukocyte Antigens. Preprint at https://www.biorxiv.org/content/10.1101/2024.04.01.587C17v2 (2024).

16. Miles, J.J., Douek, D.C. & Price, D.A. Bias in the αβ T-cell repertoire: implications for disease pathogenesis and vaccination. Immunol. Cell Biol. 89, 375–87 (2011).

17. Hallmayer, J. et al. Narcolepsy is strongly associated with the T-cell receptor alpha locus. Nat. Genet. 41, 708–11 (2009).

18. Ollila, H.M. et al. Narcolepsy risk loci outline role of T cell autoimmunity and infectious triggers in narcolepsy. Nat. Commun. 14, 2709 (2023).

19. Mikocziova, I., Greiff, V. & Sollid, L.M. Immunoglobulin germline gene variation and its impact on human disease. Genes Immun. 22, 205–217 (2021).

20. Andersen, I.L. et al. Serological screening for coeliac disease in an adult general population: the HUNT study. Gut (2025).

21. Lukina, P. et al. Coeliac disease in the Trondelag Health Study (HUNT), Norway, a population-based cohort of coeliac disease patients. BMJ Open 14, e077131 (2024).

22. Åsvold, B.O. et al. Cohort profile update: The HUNT Study, Norway. Int J Epidemiol 52, e80–e91 (2023).

23. Corcoran, M. et al. Archaic humans have contributed to large-scale variation in modern human T cell receptor genes. Immunity 56, 635–652.e6 (2023).

24. Corpas, M. et al. Genetic signature detected in T cell receptors from patients with severe COVID-19. iScience 26, 107735 (2023).

25. Posnett, D.N. et al. Level of human TCRBV3S1 (V beta 3) expression correlates with allelic polymorphism in the spacer region of the recombination signal sequence. J. Exp. Med. 179, 1707–11 (1994).

26. Moffatt, M.F., Traherne, J.A., Abecasis, G.R. & Cookson, W.O.C.M. Single nucleotide polymorphism and linkage disequilibrium within the TCR α/δ locus. Hum. Mol. Genet. 9, 1011–1019 (2000).

27. Mackelprang, R. et al. Sequence diversity, natural selection and linkage disequilibrium in the human T cell receptor alpha/delta locus. Hum. Genet. 119, 255–66 (2006).

28. Subrahmanyan, L., Eberle, M.A., Clark, A.G., Kruglyak, L. & Nickerson, D.A. Sequence variation and linkage disequilibrium in the human T-cell receptor beta (TCRB) locus. Am. J. Hum. Genet. 69, 381–95 (2001).

29. Petersen, J. et al. Diverse T cell receptor gene usage in HLA-DQ8-associated celiac disease converges into a consensus binding solution. Structure 24, 1643–1657 (2016).

30. Petersen, J. et al. Determinants of gliadin-specific T cell selection in celiac disease. J. Immunol. 194, 6112–6122 (2015).

31. Petersen, J. et al. T-cell receptor recognition of HLA-DQ2–gliadin complexes associated with celiac disease. Nat. Struct. Mol. Biol. 21, 480–488 (2014).

32. Næss, M. et al. Data resource profile: The HUNT Biobank. Int. J. Epidemiol. 53(2024).

33. Risnes, L.F. et al. Disease-driving CD4+ T cell clonotypes persist for decades in celiac disease. J. Clin. Invest. 128, 2642–2650 (2018).

34. Islam, S. et al. Characterization of the single-cell transcriptional landscape by highly multiplex RNA-seq. Genome Res. 21, 1160–7 (2011).

35. Vander Heiden, J.A. et al. pRESTO: a toolkit for processing high-throughput sequencing raw reads of lymphocyte receptor repertoires. Bioinformatics 30, 1930–2 (2014).

36. Gupta, N.T. et al. Change-O: a toolkit for analyzing large-scale B cell immunoglobulin repertoire sequencing data. Bioinformatics 31, 3356–8 (2015).

37. Ye, J., Ma, N., Madden, T.L. & Ostell, J.M. IgBLAST: an immunoglobulin variable domain sequence analysis tool. Nucleic Acids Res. 41, W34–40 (2013).

38. Manso, T. et al. IMGT® databases, related tools and web resources through three main axes of research and development. Nucleic Acids Res. 50, D1262–d1272 (2022).

39. Giudicelli, V., Chaume, D. & Lefranc, M.P. IMGT/GENE-DB: a comprehensive database for human and mouse immunoglobulin and T cell receptor genes. Nucleic Acids Res. 33, D256–61 (2005).

40. Sayers, E.W. et al. Database resources of the National Center for Biotechnology Information in 2025. Nucleic Acids Res. 53, D20–d29 (2025).

