## Supplemental Information for "A predisposing effect of HLA class II genes in celiac disease by skewing the naïve CD4^+^ T-cell receptor repertoire"

**Supplementary Table 1** Antibodies used for FACS.

| Surface marker | Fluorochrome | Manufacturer | Clone | Dilution factor |
|---|---|---|---|---|
| *Strategy 1 and 2* | | | | |
| CD3 | BV785 | BioLegend | UCHT1 | 1:33 |
| CD14 | BV510 | BioLegend | M5E2 | 1:33 |
| CD19 | PE-Cy7 | BioLegend | HIB19 | 1:25 |
| CD27 | BV421 | BioLegend | O323 | 1:33 |
| IgG* | APC | BioLegend | M1310G05 | 1:25 |
| IgA* | APC | MACS | REA1014 | 1:25 |
| CD45RB* | APC | Thermo Fisher | MEM-55 | 1:25 |
| IgD* | PE | BioLegend | IA6-2 | 1:33 |
| CD4 | APC-Cy7 | BioLegend | OKT4 | 1:50 |
| CD62L | PerCP-Cy5.5 | BioLegend | DREG-56 | 1:33 |
| CD45RA | Alexa Fluor 488 | BioLegend | HI100 | 1:33 |
| *Strategy 3* | | | | |
| CD62L | BV785 | BioLegend | DREG-56 | 1:33 |
| CD14 | BV510 | BioLegend | M5E2 | 1:33 |
| CD19 | PE-Cy7 | BioLegend | HIB19 | 1:25 |
| CD27 | BV421 | BioLegend | O323 | 1:33 |
| CD197** | APC | Miltenyi | *** | |
| CLA** | APC | Miltenyi | HECA-452 | |
| CD62L** | APC | Miltenyi | 145/15 | |
| IgD* | PE | BioLegend | IA6-2 | 1:33 |
| CD4 | APC-Cy7 | BioLegend | OKT4 | 1:50 |
| IgA* | PerCP-Vio700 | Miltenyi Biotec | IS11-8E10 | 1:50 |
| IgG* | PerCP-Vio700 | Miltenyi Biotec | IS11-3B2.2.3 | 1:50 |
| CD45RB* | PerCP-Vio700 | Miltenyi Biotec | REA119 | 1:50 |
| CD45RA | Alexa Fluor 488 | BioLegend | HI100 | 1:33 |

BV – Brilliant Violet, PE – Phycoerythrin, APC – Allophycocyanin, PerCP – Peridinin-chlorophyll-protein. * Samples were stained with these antibodies for a separate project. These markers were not used for gating in this study. ** Antibodies were not added in this stain mix, but samples had already been stained with these antibodies before cryopreservation and remained partially positive for APC after thawing. *** APC-labeled antibody added as part of the human CD4+ Effector Memory T Cell Isolation Kit.

**Supplementary Table 2** Oligo sequences used in AIRR-seq library prep of naïve T cells.

| Oligo | Step | Sequence (5'→ 3') |
|---|---|---|
| TSO | cDNA synthesis | Biotin-AAGCAGTGGTATCAACGCAGAGTNNNNNNNNNNNNCTTrGrGrG |
| Oligo d(T) | | CTGAATTCTTTTTTTTTTTTTTTT |
| STRT-fwd-Long | PCR1 | CTAATACGACTCACTATAGGGCAAGCAGTGGTATCAACGCAGAGT |
| STRT-fwd-Short | | CTAATACGACTCACTATAGGGC |
| TRAC | | GGAACTTTCTGGGCTGGGGAAGAAGGTGTCTTCTGG |
| TRBC | | TGCTTCTGATGGCTCAAACACAGCGACCT |
| R1_PCR2_TRACX | PCR2 | ACACTCTTTCCCTACACGACGCTCTTCCGATCTNNNNNNXXXXXXCAGCTGGTACACGGCAGGGT |
| R1_PCR2_TRBCX | | ACACTCTTTCCCTACACGACGCTCTTCCGATCTNNNNNNXXXXXXCGACCTCGGGTGGGAACAC |
| R2_PCR2_InX | | GGCATTCCTGCTGAACCGCTCTTCCGATCTNNNNNNXXXXXXAAGCAGTGGTATCAACGCAGAGT |
| R1 | PCR3 | AATGATACGGCGACCACCGAGATCTACACTCTTTCCCTACACGACGCTCTTCCGATC |
| R2 | | CAAGCAGAAGACGGCATACGAGATCGGTCTCGGCATTCCTGCTGAACCGCTC |

All primers were HPLC purified and from Biomers, except the TSO, which was purified by standard desalting and from IDT. Ns indicate random nucleotides, rG indicates riboguanosine, Xs indicate sample-specific indexes (see Supplementary Table 3 for individual index sequences).

**Supplementary Table 3** Index sequences used in 2nd PCR to multiplex several subjects in each sequencing library (Excel file).

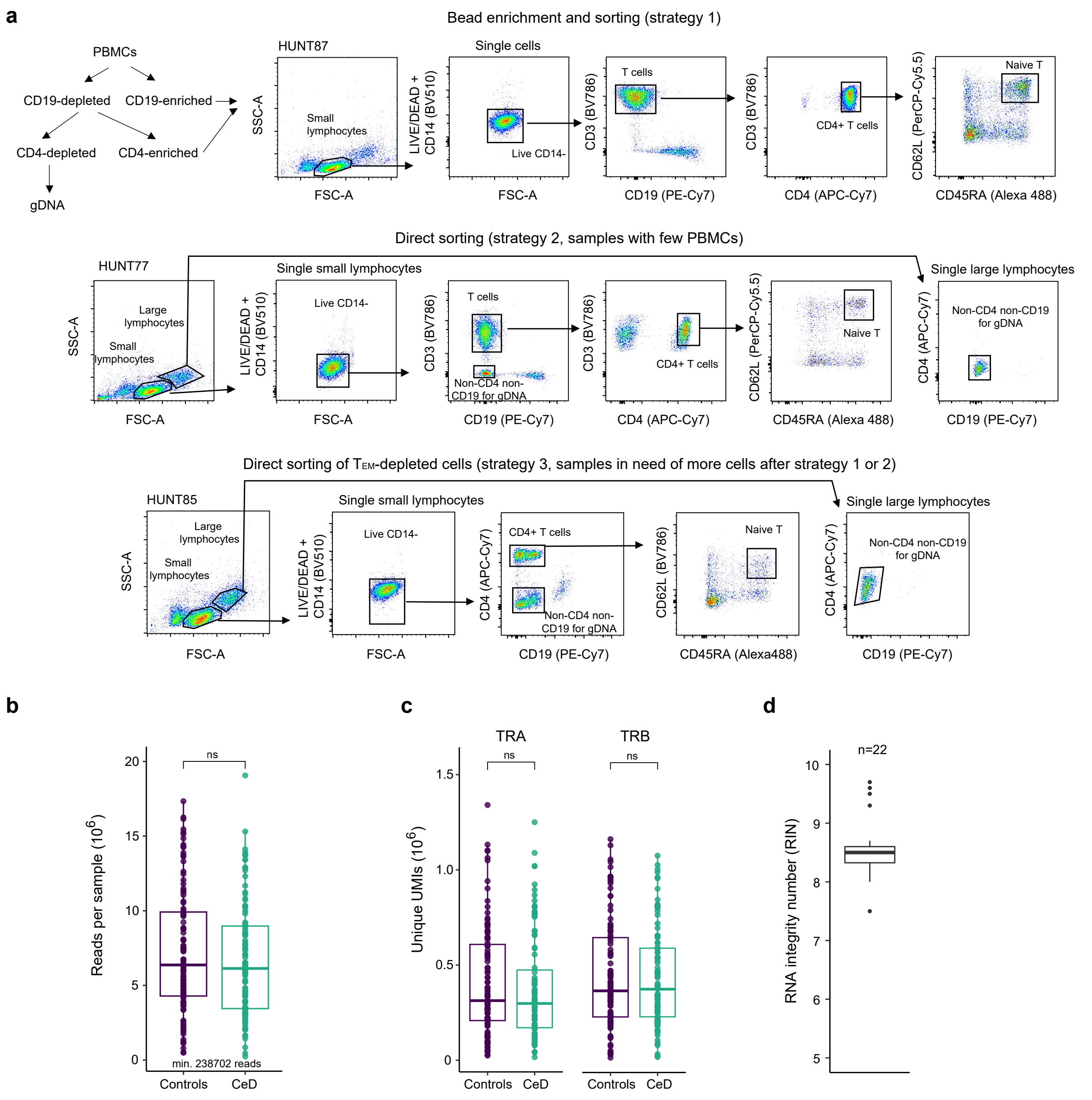


**Supplementary Figure 1** Approach and AIRR-seq quality control (related to Fig. 1) (**a**) Strategies for isolation of naïve CD4⁺ T cells and other cells for extraction of genomic DNA. (**b**) Paired R1 and R2 reads per sample after quality control. (**c**) UMIs per sample. (**d**) RNA integrity number (RIN) per tested sample, determined by Agilent Bioanalyzer.

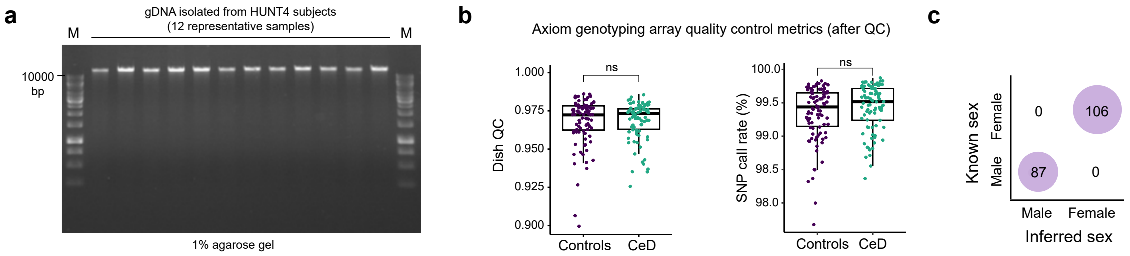


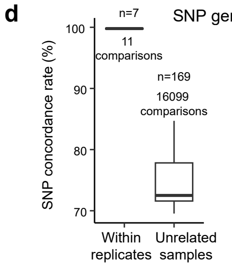


SNP genotype concordance within sample replicates and between unrelated samples

Sample replicates

| Sample | Plate | Purified | Source | Control purpose |
|---|---|---|---|---|
| HUNT117 | P2 / P2 | No | Bead depletion | Effect of gDNA concentration |
| HUNT256 | P1 / P2 / P3 | No | FACS | Plate positive control |
| HUNT187 | P1 / P2 / P3 | No | Bead depletion | Plate positive control |
| HUNT236 | P1 / P3 | No / Yes | Unsorted / FACS | gDNA purification & source |
| HUNT170 | P1 / P3 | No / Yes | Unsorted / FACS | gDNA purification & source |
| HUNT174 | P1 / P3 | No / Yes | Unsorted / FACS | gDNA purification & source |
| HUNT254 | P1 / P3 | No / Yes | Unsorted / FACS | gDNA purification & source |

**Supplementary Figure 2** SNP genotyping quality control (related to Fig. 1) (**a**) Representative examples of high integrity genomic DNA (gDNA) assessed by agarose gel electrophoresis. M indicates DNA ladder. (**b-d**) SNP genotyping of gDNA using an Axiom genotyping chip. (**b**) Sample quality metrics after quality control (QC), showing Dish QC (left) and SNP call rate (right) for control and CeD subjects. Statistical significance was calculated using a Wilcoxon rank sum test. (**c**) Known compared to computationally inferred biological sex of genotyped subjects. (**d**) Concordance analysis comparing SNP genotype calls of all samples to all other samples. Some sample replicates were included as controls (table). Comparisons between related samples (sample duplicates and triplicates) were plotted together against non-related samples to verify high within-replicate concordance and low inter-sample concordance.

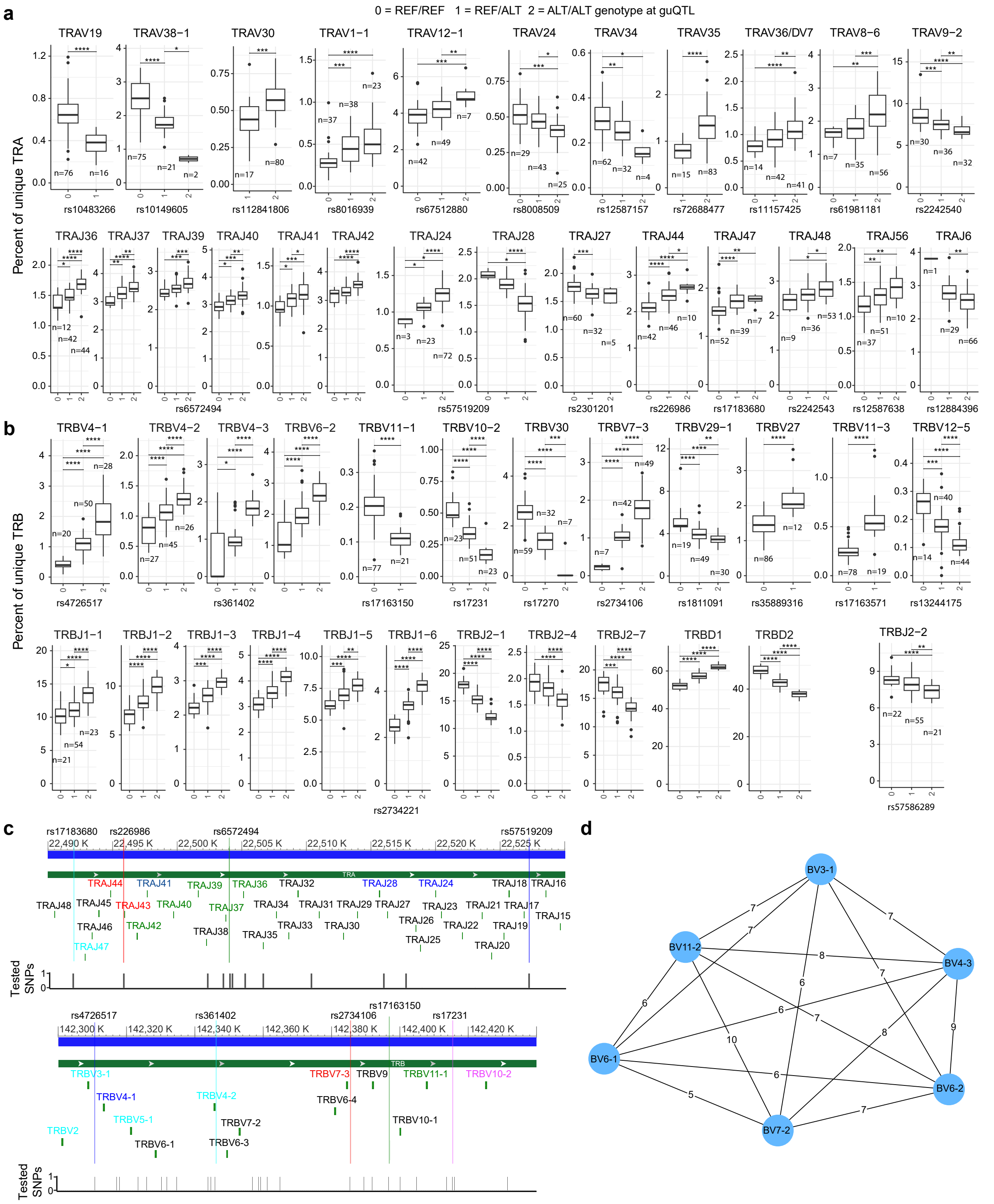


**Supplementary Figure 3** *Cis*-guQTL analysis of TRA and TRB gene usage in the naïve CD4⁺ T-cell repertoire of control subjects (related to Fig. 2). (**a,b**) Additional examples of *cis*-mediated effects of lead guQTL genotype on gene expression of TRA (**a**) and TRB (**b**) genes. Statistical significance was calculated using a Wilcoxon rank sum test for each gene and Bonferroni correction within each subfigure. (**c**) Additional examples of location of lead guQTLs with respect to their affected TRA or TRB genes. (**d**) Additional network of TRB genes associated to the same significant guQTLs.

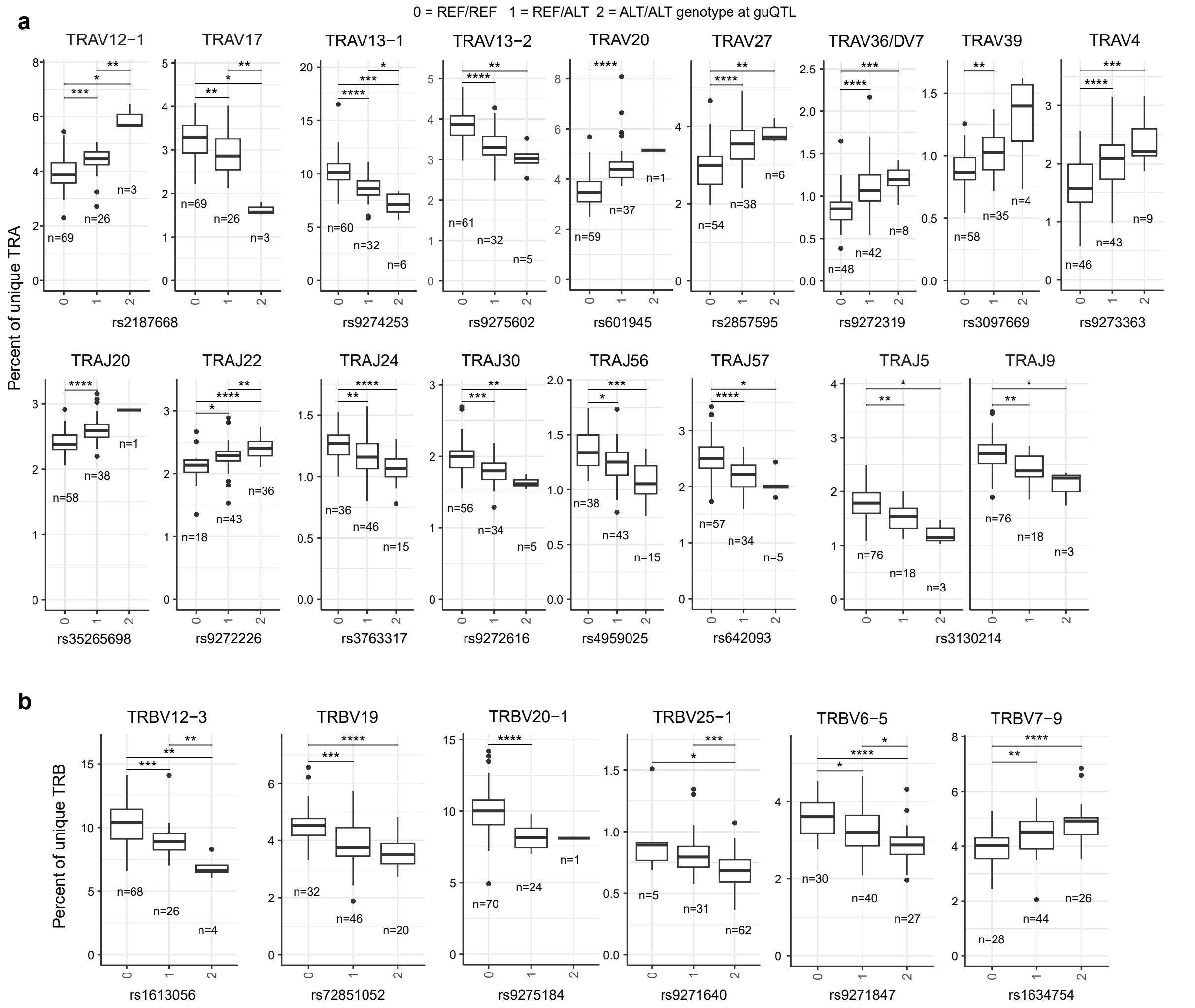


**Supplementary Figure 4** *Trans*-guQTL analysis of the influence of the HLA region on TRA and TRB gene usage in the naïve CD4+ T-cell repertoire of control subjects (related to Fig. 3) (**a,b**) Additional examples of *trans*-mediated effect of lead guQTL genotype on TRA (**a**) and TRB (**b**) gene expression. Statistical significance was calculated using a Wilcoxon rank sum test for each gene and Bonferroni correction within each subfigure. Non-significant differences are not shown.

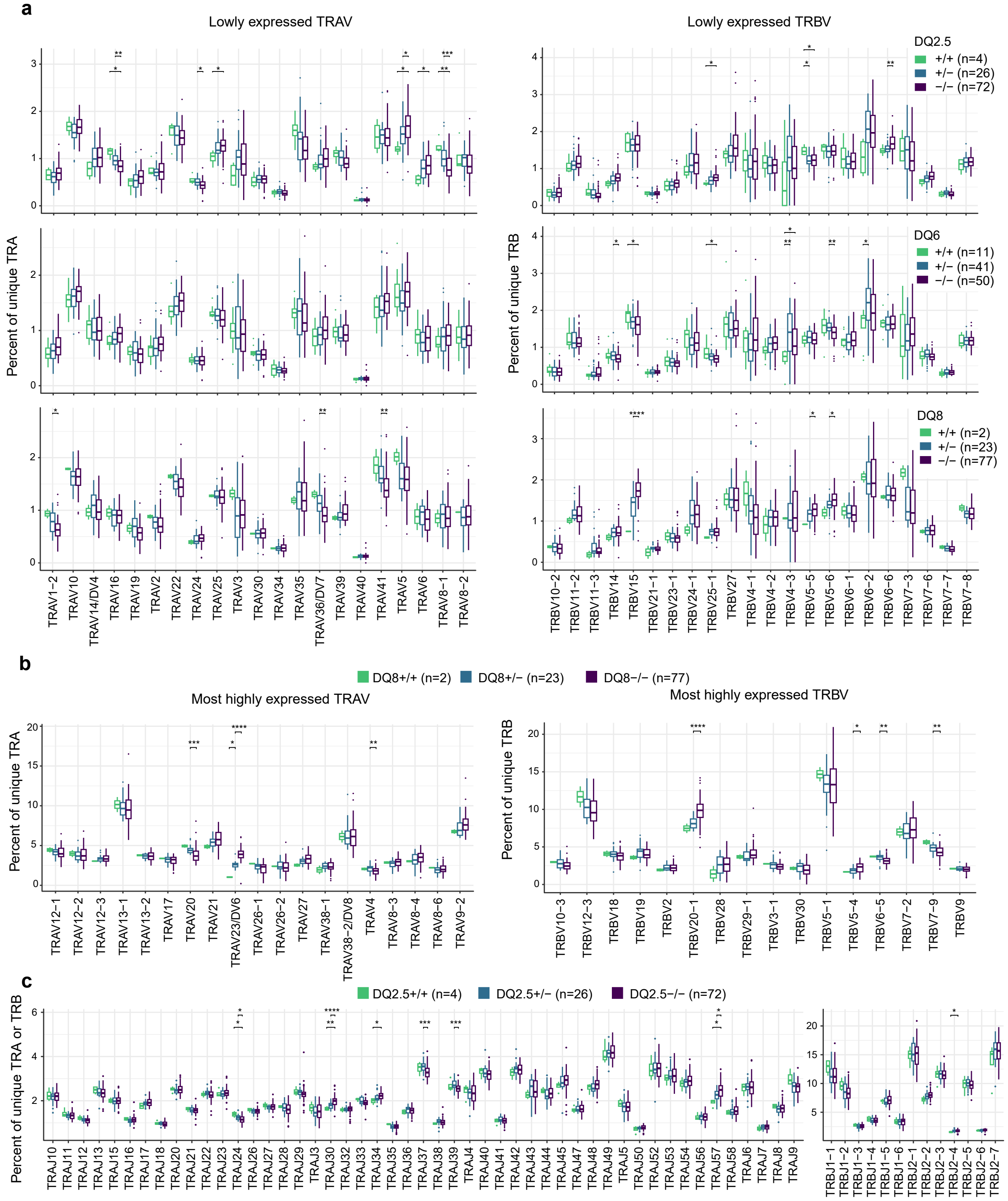


**Supplementary Figure 5** Variable TRA and TRB gene usage depending on HLA class II allotypes (related to Fig. 4). (**a**) TRAV and TRBV gene usage of control subjects depending on DQ2.5 (upper), DQ6 (middle) and DQ8 (lower) allotypes, showing lowly expressed genes. Three TRAV genes with negligible expression (median < 0.02%) and 19 TRBV genes with negligible or very low expression (median < 0.2%) were excluded. (**b**) TRAV and TRBV gene usage of control subjects depending on DQ8 allotype, showing the most highly expressed genes. (**c**) TRAJ and TRBJ gene usage of control subjects depending on DQ2.5 allotype. Nine TRAJ genes with very low expression (median < 0.2%) were excluded. Statistical significance was calculated using a Wilcoxon rank sum test for each gene and Bonferroni correction within each subfigure.

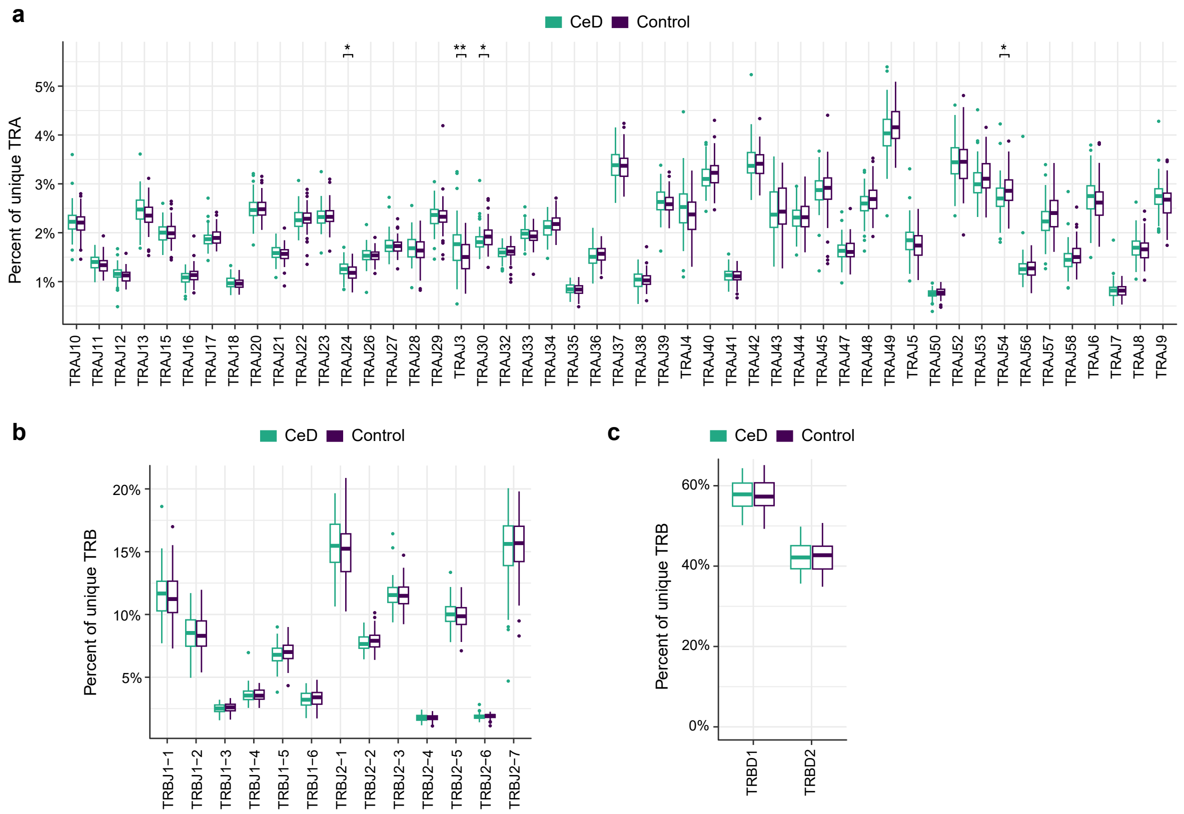


**Supplementary Figure 6** J and D gene usage distribution among naïve CD4$^+$ T cells of CeD individuals compared to controls. (**a**) TRAJ gene usage. Nine genes with very low expression (median < 0.2%) were excluded. (**b**) TRBJ gene usage. (**c**) TRBD gene usage. Statistical significance was calculated using a Wilcoxon rank sum test for each gene and Bonferroni correction within each subfigure. Non-significant differences are not shown.

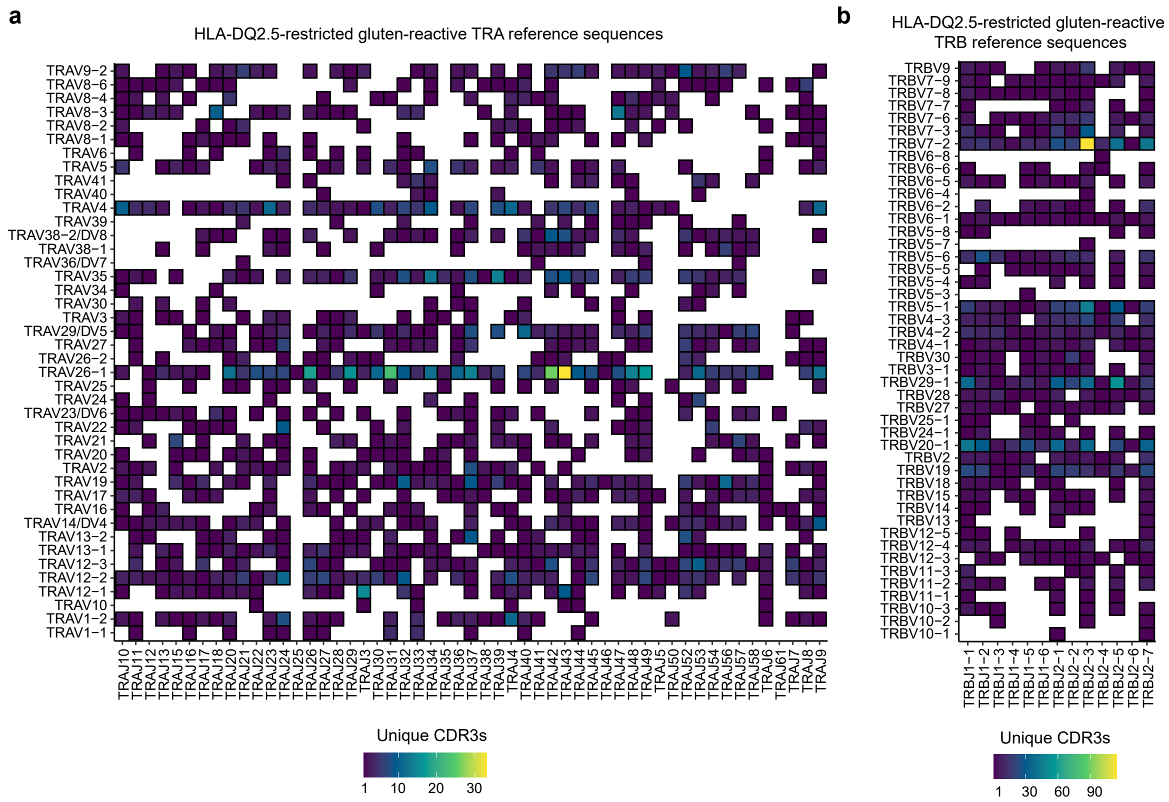


**Supplementary Figure 7** V and J gene usage distribution in a reference database of DQ2.5-restricted gluten-reactive TCR sequences from 50 CeD subjects[1], related to Fig. 6e. The number of unique TRA (**a**) and TRB (**b**) sequences (unique CDR3 region at amino acid level) for each V/J combination is indicated with color.

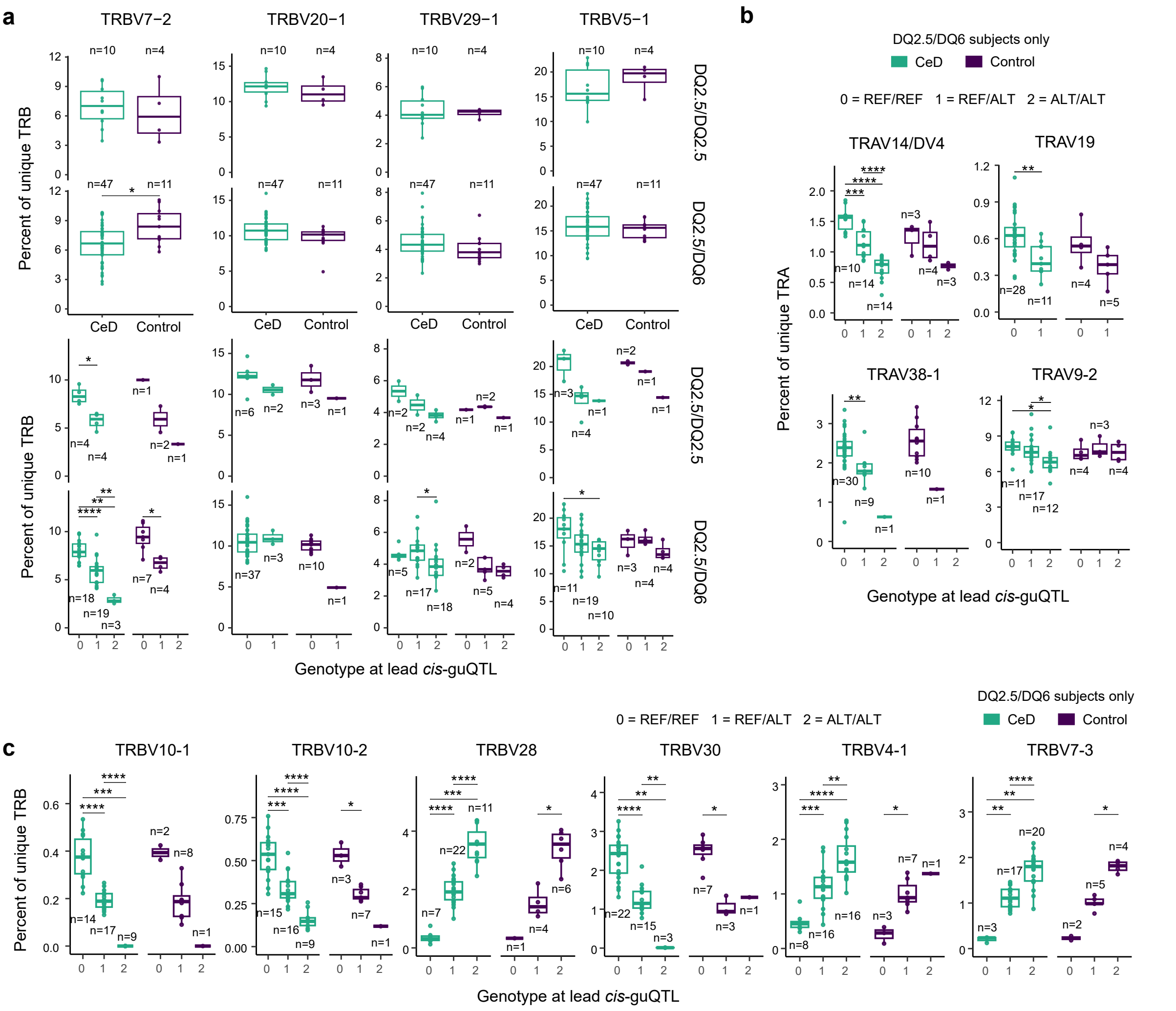


**Supplementary Figure 8** Conditional *cis*-guQTL analysis of V genes (related to Fig. 7). (**a**) Four most frequently used TRBV genes by DQ2.5-restricted gluten-reactive effector T cells in CeD. (**b**,**c**) TRAV (**b**) or TRBV (**c**) genes with most significant lead cis-guQTLs. Gene usage is conditioned by HLA-DQ types (DQ2.5/DQ2.5 and DQ2.5/DQ6 (**a**) or DQ2.5/DQ6 only (**b**,**c**)) and lead *cis*-guQTL genotype based on *cis*-guQTL analysis of all the controls. Statistical significance was calculated using a Wilcoxon rank sum test for each gene and Bonferroni correction within each subfigure. Non-significant differences are not shown.

### References

1. Dahal-Koirala, S. *et al.* Comprehensive analysis of CDR3 sequences in gluten-specific T-cell receptors reveals a dominant R-motif and several new minor motifs. *Front. Immunol.* **12**, 639672 (2021).
